# Imaging stable isotope labelling kinetics (iSILK) for following spatial Aβ plaque aggregation dynamics in evolving Alzheimer’s disease pathology

**DOI:** 10.1101/2020.10.12.335828

**Authors:** Wojciech Michno, Katie Stringer, Thomas Enzlein, Melissa K. Passarelli, Stephane Escrig, Kaj Blennow, Henrik Zetterberg, Anders Meibom, Carsten Hopf, Frances A. Edwards, Jörg Hanrieder

## Abstract

For our understanding of Alzheimer’s disease (AD) pathophysiology, it is of critical importance to determine how key pathological factors, specifically amyloid β (Aβ) plaque formation, are interconnected and implicated in neurodegeneration, disease progression and the development of clinical symptoms. Exactly how Aβ plaque formation is initiated and how the ongoing plaque deposition proceeds is not well understood. This is partly because we can only examine details of the molecular pathology after death in humans, and in mice, we can only examine a particular point in time without any longitudinal information on the fate of individually formed deposits. Herein, we used metabolic labelling of proteins with stable isotopes, together with multimodal imaging mass spectrometry (IMS) for imaging stable isotope labelling kinetics (iSILK) in the APP^NL-G-F^ knock-in mouse model of AD. The aim was to monitor the earliest seeds of Aβ deposition through ongoing plaque development and track the deposition of Aβ that is produced later in relation to already deposited plaques. This allowed us to visualize Aβ peptide aggregation dynamics within individual deposits across different brain regions. We identified the cortex as a primary site of deposition in precipitating plaque pathology. Further, our data show that structural plaque heterogeneity is associated with differential peptide deposition. Specifically, Aβ1-42 is forming an initial core seed followed by radial outgrowth and late secretion and deposition of Aβ1-38.

Together these data prove the potential of iSILK for probing amyloid protein secretion, processing and aggregation dynamics in AD pathology.

## Introduction

Alzheimer’s disease (AD) poses an immense societal challenge, particularly as there are still no disease-modifying treatments. ^1, 2, 3^ The major hallmarks of AD are the progressive accumulation of amyloid β (Aβ) peptide derived from amyloid precursor protein (APP) and intracellular deposition of hyperphosphorylated tau protein. ^4, 5, 6^ Although the importance of progressive Aβ plaque deposition in AD is long recognized, exactly how plaques develop over time and their relation to toxicity or homeostatic response of the surrounding neuronal network is not clear. ^2, 4, 7, 8^ Aβ aggregation is considered a highly dynamic process ^6^ and therefore, it is of great relevance to delineate the fate of different amyloid fibrillary structures in evolving Aβ pathology.

A major challenge in investigating Aβ pathology at submicron levels is the need for appropriate imaging technologies that combine the necessary spatial resolution, sensitivity and specificity to probe these processes. Indeed, the limited understanding of amyloidogenic protein aggregation in AD pathogenesis relates directly to the lack of effective imaging tools with high chemical, spatial and longitudinal precision. ^9, 10^

Recent advances in using metabolic, *in vivo* labelling with stable isotopes, followed by mass spectrometry were demonstrated to be a powerful tool to measure stable isotope labeling kinetics and protein turnover termed SILAC^11, 12^, SILAM^13^ or SILK^14^. In particular, in the context of AD, SILK studies have been pioneering the field of in vivo stable isotope labelling for clinical applications and specifically towards Alzheimer’s disease. In SILK, intravenous infusions of ^13^C6-Leu, allowed the quantification of Aβ and tau turnover dynamics in continuously collected CSF from living AD patients, revealing significant insights into protein clearance impairment associated with plaque pathology. ^14, 15^ While these tools are of significant importance, they provide limited information spatial information on isotope incorporation.

Novel chemical imaging technologies such as imaging mass spectrometry ^16^ greatly increase the resolution of such events. The combination of isotope labelling of proteins with nanoscale secondary ion imaging (NanoSIMS) of such isotopes, termed multi ion beam imaging (MIMS), has made it possible for measuring spatial protein turnover kinetics in cells and tissues, and most significantly demonstrated isotope incorporation in plaques in a hospice study of AD patients as well as in transgenic APP mice, also referred to as SILK-SIMS. ^17, 18^

One challenge with SIMS imaging is the limited molecular information retrievable, permitting analysis of intact peptides and proteins. This is something that can be achieved with matrix assisted laser desorption ionization mass spectrometry based IMS (MALDI-IMS), which has been successfully demonstrated for monitoring spatial metabolization of i.p. injected, labeled drug candidates. ^19^ Most significantly, MALDI-IMS allows for chemically specific Aβ peptide imaging of Aβ pathology in animal models of AD ^10, 20^ and in post mortem human AD brain tissue. ^21, 22^

We here set out to take advantage of these novel isotope labelling and imaging methods to follow Aβ peptide specific aggregation dynamics in a recently developed knock-in mouse model of AD with familial mutations in amyloid precursor protein (APP; APP ^NL-G-F^). ^23^

We performed metabolic labelling with stable isotopes of these mice combined with multimodal imaging mass spectrometry, including MALDI-IMS and NanoSIMS, for spatial delineation of differential stable isotope label incorporation in situ. Given that this imaging paradigm are implemented and applied in the context of AD pathology this approach is herein hence referred to as imaging SILK (iSILK). For the first time, the here described iSILK experiments allowed for visualizing aggregation dynamics of different Aβ peptides within single plaques and across different brain regions in evolving plaque pathology from early deposition to later plaque growth. Specifically, by using a comprehensive labelling scheme, we found that Aβ pathology in APP ^NL-G-F^ mice precipitates in the cortex by forming small dense-core deposits consisting of Aβ1-42. Later events in early plaque pathology involve plaque growth upon homogenous Aβ1-42 deposition, deposition in the hippocampus, and secretion and deposition of Aβ1-38.

## Results

### 1. Stable isotopes can be metabolically incorporated in specific Aβ peptides and accumulate into extracellular plaques upon amyloid pathology onset

A central issue in delineating AD pathogenesis is that amyloid plaque pathology starts to develop long before any cognitive symptoms occur. It is therefore of great relevance to delineate the fate of different amyloid fibrillary structures in evolving Aβ pathology, from early accumulation and aggregation, to deposition and maturation of extracellular plaques.

However, this is challenging, as early accumulation, aggregation and deposition of Aβ peptides are highly dynamic processes. We therefore set out to examine the Aβ aggregation dynamics in APP^NL-G-F^ mice. We aimed to delineate the initial events and sites of plaque onset in the brain by following amyloid aggregation using an iSILK imaging paradigm. This was based on metabolic labelling with stable isotopes, followed by spatial delineation of isotope-resolved aggregation-kinetics using multimodal nanoSIMS and MALDI-IMS.

We provided ^15^N-labelled protein diet to APP^NL-G-F^ mice from age 7-17 weeks (11-week PULSE period, Fig. 1A) to ensure that all APP is labelled prior and during initial Aβ secretion, Aβ deposition and plaque onset, which in the APP ^NL-G-F^ mouse model takes place at around week 8-9. ^23^ Consequently, all the Aβ that was secreted and accumulated during rising amyloid just prior to deposition should contain ^15^N label. A CHASE period was omitted, to avoid interference of heterogeneously labelled Aβ produced during a potential washout.

**Figure 1:**
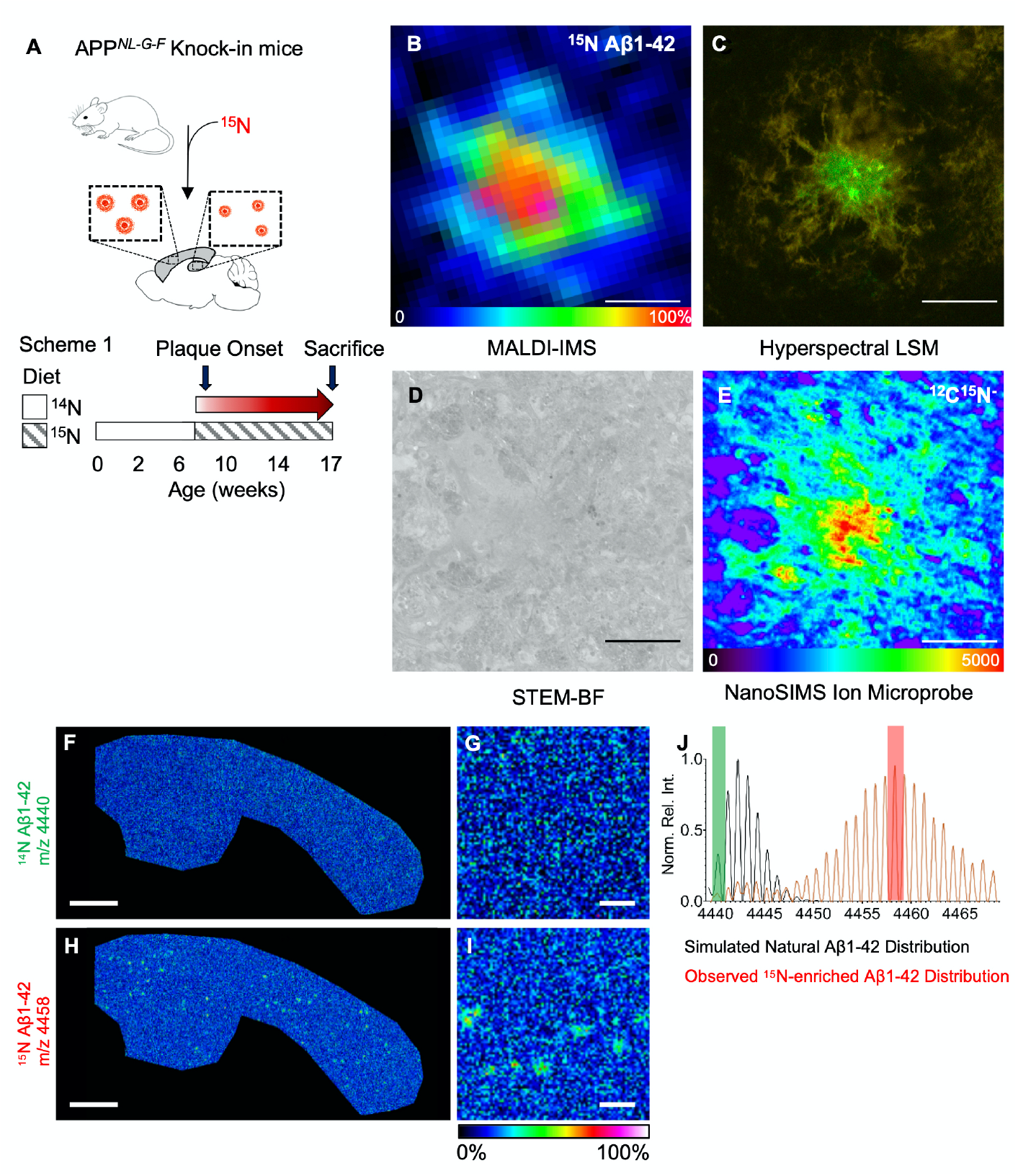
iSILK enables analysis of heavy-isotope incorporation at a single amyloid-β (Aβ) plaque level. **(A)** Labeling scheme to establish feasibility of ^15^N incorporation at the onset of Aβ plaque pathology in knock-in APP^NL-G-F^ mice, and reveals inter- and intra-plaque heterogeneity. Sample images of individual Aβ plaques obtained through four distinct imaging modalities, including: **(B)** MALDI-IMS total ion image of labelled Aβ1-42 (m/z 4450-4466); **(C)** Hyperspectral laser-scanning confocal microscopy (Hyperspectral LSM;); **(D)** Annular bright-field imaging using scanning transmission electron microscopy (STEM-BF) and **(E)** NanoSIMS Ion Microprobe Imaging. (E) Int scale: [^15^N^12^C^-^] m/z 27. Scalebar B-E: 10 μm **(F-I)** MALDI-IMS of Aβ peptide signals following 11 weeks ^15^N diet, starting week 7. **(F,G)** No unlabeled Aβ peptides were detected these mice and only Aβ species containing ^15^N were observed that show distinct accumulations **(H-I)** that were LCO positive **(C)**. The degree of ^15^N incorporation is further visualized in MALDI-MS spectra acquired in reflector positive mode from LMD-IP extracted plaques **(J)**. Scalebar F,I 1000 μm; G,J: 75 μm

We analyzed the stable isotope enrichment in brain tissue and plaques in these mice using multimodal chemical imaging (Supplementary Information Fig. S1), including MALDI-IMS (Fig. 1B), fluorescent structural amyloid staining (Fig. 1C), scanning transmission electron microscopy (STEM, Fig. 1D) and nanoscale SIMS (Fig. 1E). Further, we validated the Aβ identity of the deposits detected with MALDI-IMS and luminescent conjugated oligothiophene (LCO) microscopy using immunohistochemistry (Supplementary Information Fig. S2) as well as by off-tissue analysis using laser microdissection (LMD) together with immunoprecipitation and mass spectrometry (Supplementary Information Fig. S1-4) ^21^. This further allowed Aβ sequence validation of the intact peptide signals detected in MALDI-IMS (Supplementary Information Fig. S2-4)

The results show that ^15^N-labelled amino acids are incorporated into APP and Aβ, and deposit within Aβ plaques upon plaque formation (Fig. 1B-E). In detail, MALDI-IMS detected solely labelled ^15^N-Aβ1-42, but no unlabeled ^14^N-Aβ1-42, demonstrating that PULSE feeding began early enough to capture all of the initial secretion and deposition of Aβ.

Here ^15^N-Aβ1-42 showed distinct localization to amyloid plaques (Fig.1B). Hyperspectral microscopy of these plaques using fluorescent structure specific amyloid staining with LCO (Fig 1C) ^24^ revealed the core structure to correspond to denser Aβ aggregates (Supplementary Information Fig. S2). Further STEM imaging showed that clear fibrillary core structures were present in the center of these Aβ plaques (Fig. 1D, Supplementary Information Fig. S5,6). NanoSIMS of these plaques revealed distinct plaque structures that showed characteristic enrichment of ^15^N label with pronounced deposition at the core, as revealed by ^12^C^15^N-signal (m/z 27) (Fig. 1E), while no label was detected in control animals (Supplementary Information Fig. S6).

Further, the MALDI-IMS peptide data show that plaques in APP^NL-G-F^ mice at 17 weeks contain predominantly Aβ1-42 along with Aβ1-38. The plaque-associated Aβ species observed at the early age, 17 weeks, were found to contain solely ^15^N-labeled Aβ peptides and no unlabeled ^14^N-Aβ (Fig. 1F-J). The absence of unlabeled Aβ peptides in plaques of these labelled animals verifies that starting labelling at week <7 allows for in time, metabolic introduction of ^15^N to cover the plaque onset period from before Aβ plaque deposition.

These results show that the general iSILK labelling and analysis paradigm proved suitable to generate metabolically labelled Aβ that deposited in extracellular Aβ plaques. The data show that initiation of ^15^N feeding right before the onset of Aβ plaque pathology, is sufficient to achieve ^15^N labelling of all the Aβ peptides present in extracellular Aβ plaques. Moreover, complementary structural imaging using both fluorescent (LCO) and electron microscopy (STEM) identified formation of morphological heterogenous Aβ plaques (Fig. 2A-G). This allows for correlation of changes in labelling degrees of individual Aβ peptides to plaque polymorphism and what intra-feature structures evolve in what order during plaque formation.

**Figure 2:**
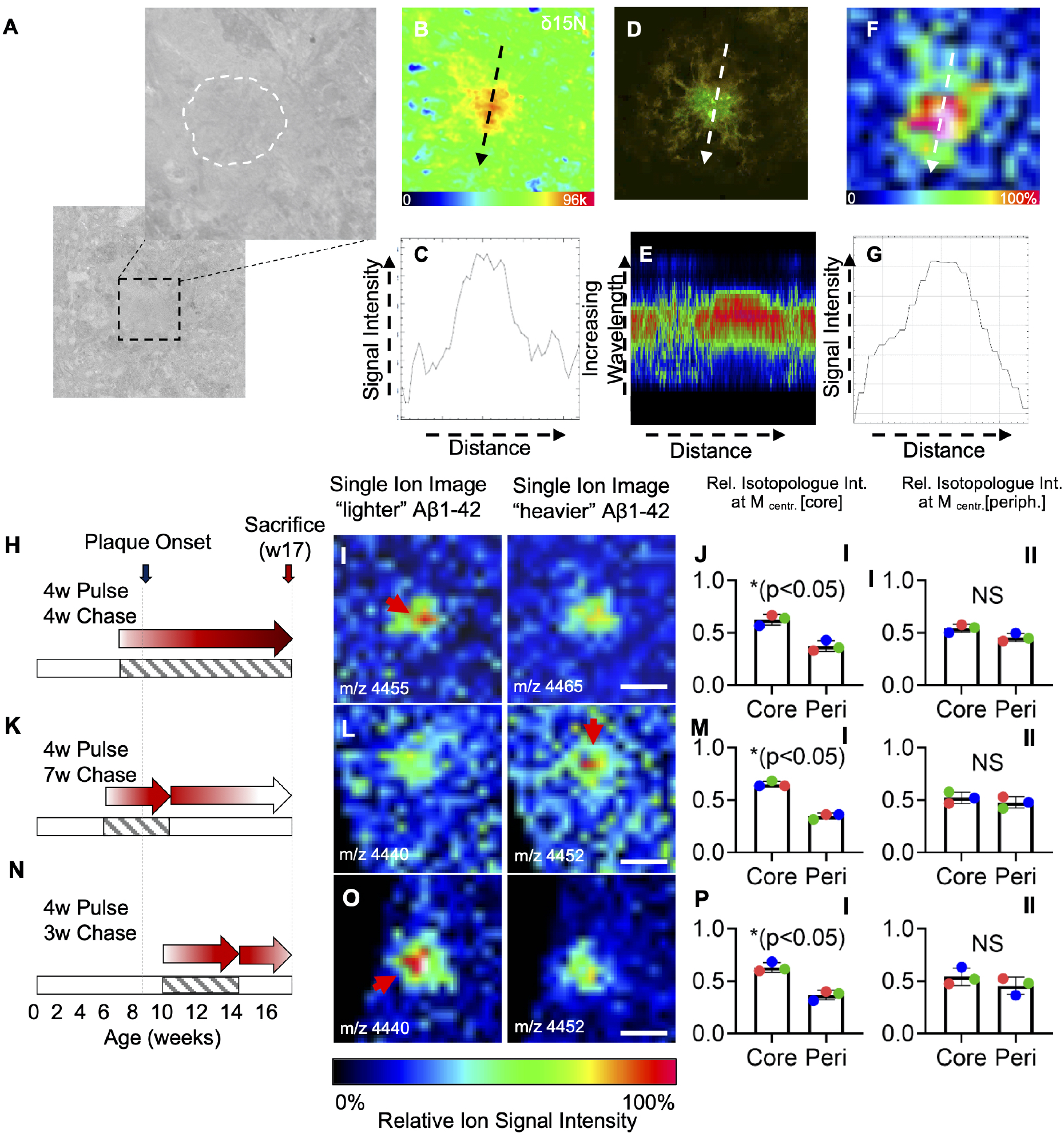
Intra-plaque heterogeneity identifies core formation to precede Aβ plaque growth. **(A-G)** Multimodal, structural imaging of plaque heterogeneity. **(A)** Sample STEM-BF image of a plaque reveals presence of a fibrillary core-like structure (zoom -outlined). **(B)** ^15^N/^14^N NanoSIMS ratio image confirming differential isotope composition of the plaque center, with higher ^15^N/^14^N signal ratio as revealed by line profile (arrow) shown in **(C)**. **(D)** Hyperspectral LSM reveal higher degree of Aβ aggregation at the core structure, as indicated by spectral shift towards shorter wavelengths in the center of the plaques trough cross-sectional spectral analysis **(E**). **(F)** MALDI-IMS total ion image of ^15^N Aβ1-42 (m/z 4450-4465) show stronger total ^15^N-Aβ1-42 signal at the center of the Aβ plaques revealed through line profile analysis **(G)**. **(H-P)** Differential labelling schemes identify, time resolved plaque formation dynamics. **(H-J)** Scheme 1: 11-week PULSE, no CHASE; **(K-M)** Scheme 2: 4-week PULSE (week 6-10), 7-week CHASE. **(N-P)** Scheme 3: 4-week PULSE (week 10-14), 3-week CHASE; Bar plots J,M,P indicate mean ± SD, MALDI Intensity scale F,I,LO: 0-100% rel. ion int.; Scalebars I,L,O: 30 μm

### 2. Initial plaque formation in APP^NL-G-F^ mice starts with formation of a dense core

AD pathology presents itself with a wide phenotypic heterogeneity among AD patients, including the formation of morphological heterogeneous plaques, including diffuse and cored deposits. ^5, 25, 26^ As diffuse plaques are also observed in amyloid positive individuals without cognitive defects ^21, 27^, Aβ deposition into cored plaques has been linked with neurotoxic pathology in a wide variety of familial AD cases, as well as in sporadic AD. ^25, 28, 29^ The actual chain of events underlying heterogenous plaque pathology and formation of diffuse and cored plaques, respectively, are still not fully understood.

This can be addressed using iSILK, where spatial mapping of relative isotope accumulation across histological features allows delineation of the degree of Aβ deposition over time within different plaque structures.

A major observation for the first experiments showed differences in the degree of labelling of the Aβ peptides across the plaque structure. MALDI-IMS showed that the total ^15^N-Aβ1-42 signal was highest in the center of Aβ plaques (Fig. 1B). Similarly, the corresponding STEM and NanoSIMS data (Fig. 2A,B) revealed a much higher ^15^N/^14^N signal to be associated with these core structures as demonstrated by the line profile for NanoSIMS (Fig. 2C). Core-specific ^15^N accumulation was further observed for plaques analyzed by complementary LCO staining (Fig. 2D) and MALDI-IMS (Fig. 2E), where a blueshift of the emission spectra across the plaque outlines the core structure (Fig. 2F) that correlates with increased ^15^N Aβ1-42 MALDI signal, as indicated in the corresponding line scan (Fig. 2G).

When considering the total relative ^15^N signal and that the degree of labelled peptide corresponds to the degree of deposition, this may have two different explanations. On the one hand, more labeled peptide results from longer, homogenous deposition period and hence earlier deposition at the core. On the other hand, more isotope signal could indicate merely a more core-specific peptide deposition in general that can take place at any time during the labelling period. This highlights the need for using complementary techniques with specificity towards the distinct Aβ peptides, something warranted by MALDI-IMS.

Indeed, inspection of the MALDI-IMS peptide spectra for single plaque ROI shows a characteristic isotope envelope for ^15^N-labelled Aβ species reflecting the degree of stable isotope incorporation and deposition (Fig. 1J). The generation of single ion images for various m/z species within the isotope envelope for Aβ peptides, allows spatial delineation of deposition patterns of differentially labelled isoforms of the same Aβ peptide (Supplementary Information Fig. S7A).

By monitoring Aβ1-42 deposition in the first labeling experiment (Fig. 2H), we observed that less labeled ^15^N-Aβ1-42 species localized to the core as compared to the more peripheral parts of the plaque (Fig. 2I, J-*I*), while more label containing ^15^N-Aβ1-42 species showed no significant localization to either the core or periphery, and deposited evenly over the plaque region (Fig. 2I, J-*II*). Since, the less labelled ^15^N-Aβ1-42 must originate from earlier secreted, and likely earlier deposited Aβ1-42, deposition at the core suggests that core structure formation is an initial event in seeding plaque pathology. The deposition pattern of later synthesized Aβ1-42, as reflected in increased content of ^15^N, is more homogenous.

Together these data point towards plaque formation in APP^NL-G-F^ being initiated by seeding as dense core and outward growth as a consequence of later, plaque wide deposition across this initial seeded plaque structures. This thus results in morphological heterogenous cored deposits with a diffuse corona.

The data from the first 11week full stop labelling scheme (Scheme 1, Fig. 2H-J), without CHASE period, suggested that increased isotope deposition at the core of the plaque is a consequence of more extended deposition. We therefore hypothesized that plaque formation in APP^NL-G-F^ mice is initiated by seeding as a dense core, and plaque growth is a consequence of later, more homogenous deposition across seeded plaque structures.

To address this question, we performed a number of PULSE/CHASE labelling schemes (Fig. 2K-P) to test and validate the findings on plaque formation observed for Scheme 1. Further, we designed shorter labelling schemes that include a CHASE/washout period allowing for introduction of unlabeled Aβ, in order to further delineate consequential events within a shorter timescale upon a certain stimulus.

To this end, we designed two more PULSE/CHASE schemes involving a 4-week ^15^N PULSE starting at either week 6, before plaque deposition (Scheme 2, Fig. 2K-J), or week 10 during plaque deposition (Scheme 3, Fig. 2N-P).

For Scheme 2, we supplied a ^15^N diet from week 6 to 10 followed by a 7-week CHASE. The aim was to ensure ^15^N labelling of APP at an even earlier stage prior to plaque pathology onset, as determined in Scheme 1. Using a shorter PULSE and a CHASE period allowed us to delineate the fate of early synthesized Aβ as compared to Aβ secreted and deposited during the washout. This can be achieved by mapping spatial intensity patterns of highly labelled Aβ species in comparison to their less and unlabeled isoforms as encoded in their isotope envelope pattern.

Our results show that the maximally labeled Aβ ^15^N-1-42 species (containing max. 9 ^15^N) was localized to the plaque core (Fig. 2L, M-I). Conversely, unlabeled Aβ1-42, which was consequently produced after washout, was found to show an even distribution pattern across the plaque area (Fig. 2L, M-I).

This is in line with the results obtained for Scheme 1, where early, less labelled Aβ ^15^N-1-42 localized to the core while late, more labelled Aβ ^15^N-1-42 localized homogenously across the plaque. Together, this further strongly suggests that dense core formation represents the earliest seeding event in extracellular plaque deposition in the APP^NL-G-F^ mice used in this study. This can be attributed to the arctic mutation of APP leading to increased hydrophobicity of full-length Aβ 1-42_arc_, and is well in line with immunohistochemical observations of young App^NL-G-F^ mice ^23^ (Supplementary Information Fig. S8) as well as transgenic APP mice carrying the same mutations (tgArcSwe). ^23^

To further validate these results, we performed an additional labeling scheme (Scheme 3, Fig. 2N-P), where ^15^N diet was supplied at a later time than in Schemes 1 and 2 (week 10 instead of week 6, Fig. 2N). The aim was to dissect the initial plaque formation phase (week 8-10) from the following phase of early plaque growth (week 10 onwards). We hypothesized that early produced Aβ will not contain any label and will deposit specifically at the core during pathology onset, similar to the results obtained for Schemes 1 and 2 (Fig. 2H,K).

Indeed, the MALDI-IMS data for Scheme 3 shows a distinct, predominant localization of unlabeled Aβ1-42 to the core in cortical deposits (Fig. 2M). Conversely, later synthesized, labelled ^15^N-Aβ1-42 shows a homogenous distribution pattern across the plaque ROI (Fig. 2M). This further supports the findings of the previous schemes and verifies that Aβ pathology in App^NL-G-F^ mice appears to be initiated through formation of small core seeds followed by homogenous plaque-wide deposition during plaque growth (Supplemental Information Fig. S8) ^23^.

### 3. Plaque pathology in APP^NL-G-F^ evolves through initial deposition of Aβ1-42 in the cortex, followed by deposition in the hippocampus

Along with intra-plaque specific, structural details on plaque growth in APP^NL-G-F^ mice, iSILK allows us to identify deposition dynamics across different brain areas. This is of relevance to identify how plaque pathology spreads across the brain and ultimately, to resolve the chain of mechanisms in how plaque formation affects vulnerable brain regions leading up to cognitive defects.

The aim was therefore to quantify the relative amounts of plaque-associated Aβ1-42 across the cortical and hippocampal deposits. We hypothesized that the deposited Aβ1-42 species that constitute the earliest isoforms for plaques in specific brain regions allow for relative comparison of where across these brain regions plaque pathology is precipitating first.

For this we isolated plaque ROI spectral data of hippocampal and cortical deposits and compared the isotopologue signature of plaque-associated Aβ1-42 that encodes the degree of labelling, as indicated by the shift in m/z value. Based on the labelling scheme, this shift in the isotopologue signature indicates earlier or later deposition. For unbiased, quantitative comparison of label incorporation, we performed curve analytics of the isotope envelope for plaque-associated Aβ1-42 signal that in turn encodes the degree of label incorporation. Here, to estimate the shift in m/z value as caused by the ^15^N label and hence difference in label content, we compared the centroid of the isotope envelope (Fig 3A).

**Figure 3:**
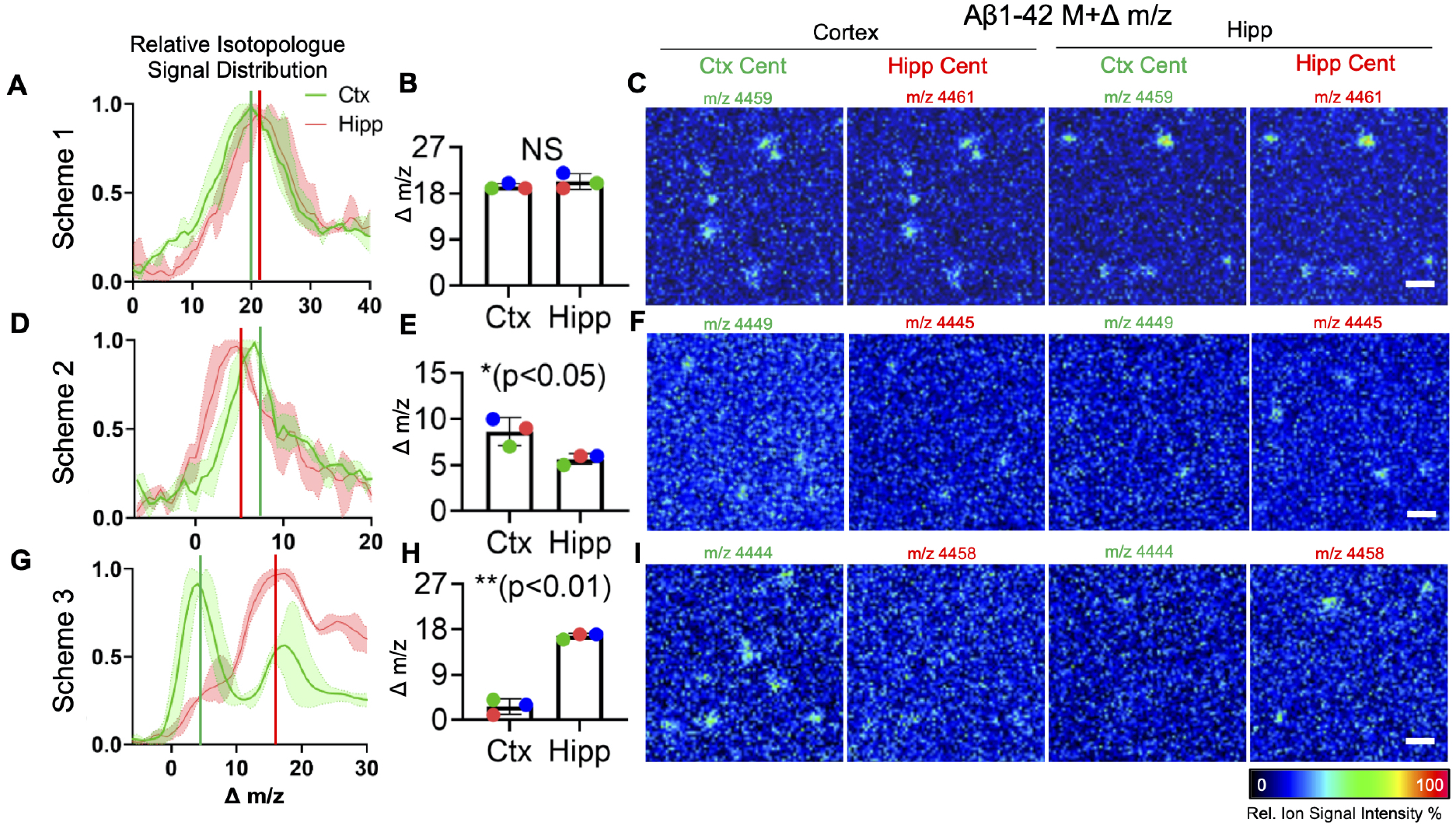
Single Aβ plaque quantification identifies primary regions for plaque deposition, and key Aβ peptide species responsible for this process. Comparative analysis of MALDI-IMS spectral data of plaque ROI for Aβ1-42 isotopologue distributions **(A)** and statistics on the centroid values **(B)** for Scheme 1. The data show no difference between cortical and hippocampal plaques. **(C)** Single ion images of Aβ1-42 in each region. The visualized Aβ1-42 signal corresponds to the different centroid m/z values for the corresponding Aβ1-42 isotope distributions observed for plaques in the Ctx vs Hipp. as a measure of average of label incorporation. Due to the overlap of isotopologue distributions, no difference is observed for Scheme 1. **(D,E)** For Scheme 2, comparison of signal from individual plaques in cortex and hippocampus, revealed higher ^15^N Aβ1-42 signal being present in cortical plaques. **(F)** Single ion maps for Scheme 2 show that though better separated, both isotope distributions are still overlapping at their respective centroids. **(G,H)** For Scheme 3, cortical plaques showed significantly less isotope incorporation (75% less) as comared to hippocampal plaques. **(I)** Single ion images verify that no low labelled (early) Aβ1-42 is detected in the Hippocampus and conversely no higher labelled Aβ1-42 is detected in the cortex. Scalebar: C,F,I: 75 μm.

For Scheme 1, Aβ1-42 no difference in label incorporation as compared to hippocampal plaques (Fig. 3A,B) attributed to the large overlap of the two distributions as further highlighted in the single ion images that do not allow to distinguish plaques with earlier and later labelled Aβ 1-42 (Fig. 3C). However, NanoSIMS imaging of cortical and hippocampal plaques showed a significantly higher degree of total ^15^N isotope incorporation of plaques in the cortex as compared to the hippocampus (Supplementary Information Fig. S9). This would suggest that plaques in the cortex are formed earlier and can thereby accumulate more ^15^N over time.

Initial deposition of plaques in the cortex is further supported by the results obtained for Scheme 2. Here, using a short PULSE and inclusion of a CHASE period showed even more pronounced differences in regional label content. In this experiment, Aβ1-42 in cortical plaques showed a 1.75-fold increase in label incorporation (175%, p<0.05) as compared to hippocampal plaques (Fig. 3D,E). This is in line with the assumption that in this labelling scheme, which started before plaque onset, early deposited Aβ1-42 contains more ^15^N label as compared to later Aβ1-42 deposited produced during the washout phase. This is further illustrated in the single ion maps generated for the different centroid m/z values (Fig. 3F), though discrimination is complicated as those ion maps show distribution of isotopologues that represent average incorporation values and show a large overlap at the corresponding m/z value (Fig. 3D and 3F). This is due to the isotope distribution pattern of different regional plaque types showing too much overlap. However, as the statistical analysis provide an unbiased overview of differences in average label incorporation, visualizing the maximum m/z values of the corresponding isotopologue distributions can illustrate exclusive deposition patterns of those very early secreted and deposited peptides. Indeed, single ion maps for maximal Aβ1-42 isotopologues show solely deposition in the cortex but not hippocampus (Supplemental Information Fig. S10) indicating that the cortex is the initial site of plaque deposition in this mouse model.

Finally, by using a complimentary labelling Scheme 3, starting after plaque onset, we aimed to further verify these results and even achieve a pronounced separation of the differentially labelled species. In this scheme, early formed plaques should contain greater amounts of early, entirely unlabeled Aβ1-42 leading to more distinct isotopologue patterns in between the regions. Indeed, for Scheme 3, Aβ1-42 in cortical plaques showed 4-fold less label incorporation (25%, p<0.01) as compared to hippocampal plaques (Fig. 3G,H). Single ion maps of the corresponding centroid m/z values, representing average isotope incorporation of the different plaque populations, show distinct differences in spatial deposition pattern. Here, low labelled, early, Aβ1-42 peptides are solely observed in the cortex, while later secreted more labelled Aβ1-42 peptides were exclusively detected in the hippocampus.

Taken together, these data show that cortical plaques form earlier than hippocampal deposits, which is in agreement with previous data described for both this mouse model as well as other, transgenic mice carrying the arctic and Swedish APP mutation. ^23, 30, 31^

### 4. Formation and deposition of C-terminally truncated Aβ1-38 occurs after initial plaque formation

The initiation and progression of amyloid plaque pathology has been suggested to be associated with secretion and aggregation of distinct Aβ peptide truncations, where both C- and N-terminal truncations have been reported. ^28, 29, 32, 33^

Given the young age of the mice used in this study, this posed an opportunity to investigate what other terminally truncated species are associated with the initial Aβ pathology in the APP^NL-G-F^ mice. The MALDI-IMS data for all labeling schemes revealed presence of Aβ1-38 in plaques, however at significantly lower amounts than Aβ1-42. This suggests that C-terminal processing and deposition are later events in early, developing plaque pathology. However, it is still under debate where and when formation and deposition of different Aβ processing products take place. We hypothesized that iSILK can be used to delineate the chain of events during sequential Aβ production and deposition; i.e. whether Aβ1-38 production occurs either through processing of early secreted, full length Aβ1-42 but through delayed deposition, or if this Aβ1-38 secretion and deposition are time-linked processes that take place later on.

We therefore compared the degree of labelling as well as the spatial deposition pattern of differentially labelled Aβ1-38 isoforms across the different labeling schemes. We further performed additional labeling schemes, deduced from Scheme 3 but with varying CHASE lengths to further delineate and validate the fate of individual labelled, Aβ1-38 isoforms over time.

For Scheme 1, we observed that the normalized degree of ^15^N labelling for Aβ1-38 deduced from the centroid values of the isotope distribution curve was not statistically different from that of Aβ1-42 (Fig. 4A,B). Further, the differentially labelled Aβ1-38 isoforms, including both less and more labelled ^15^N species, displayed a similar spatial distribution pattern across the plaque (Fig. 4C).

**Figure 4.**
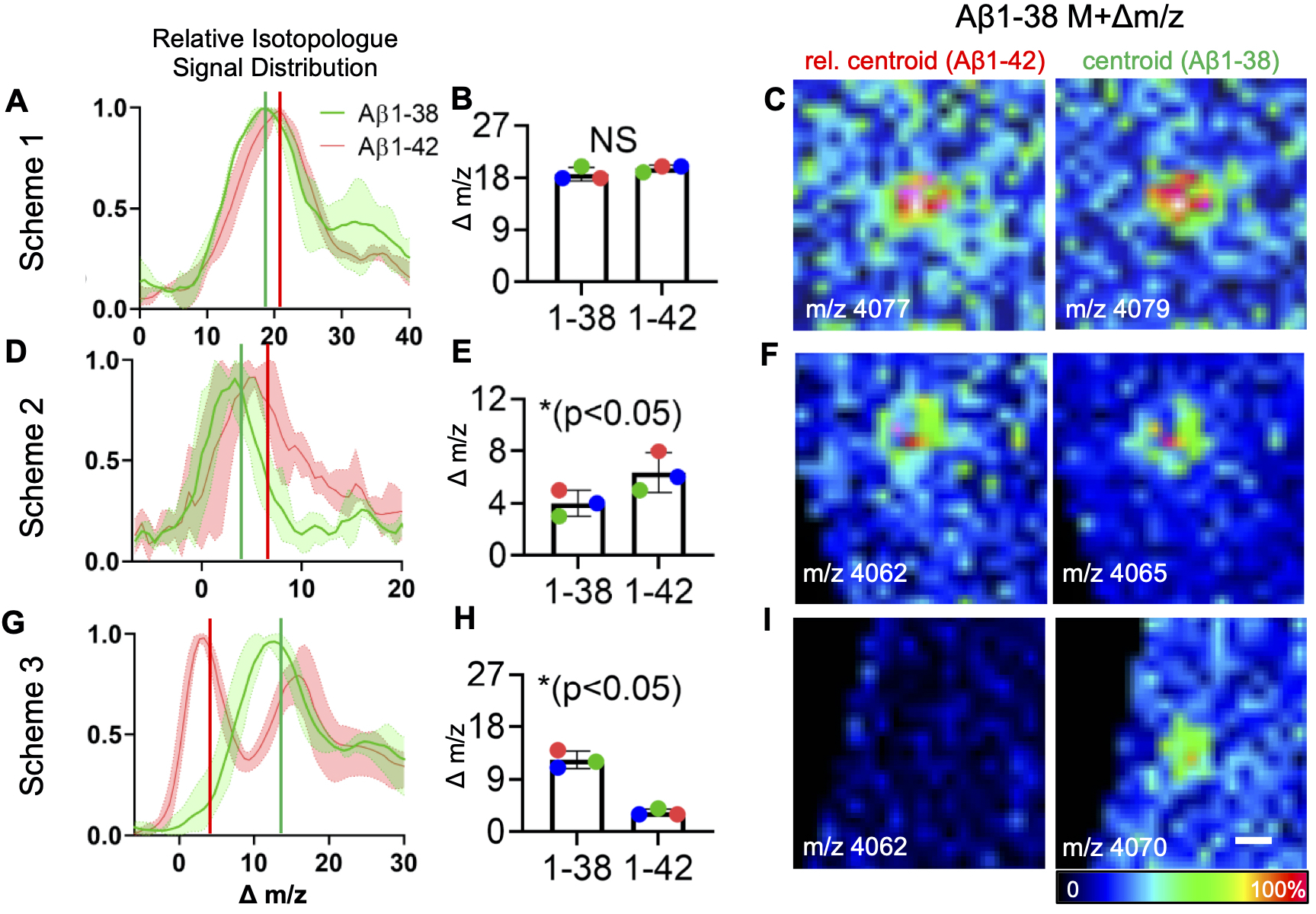
Deposition of Aβ1-38 peptide occurs after Aβ1-42 as indicated by differences in extent of ^15^N incorporation. **(A**) Comparative analysis of MALDI-IMS spectral data of plaque ROI for Aβ1-42 and Aβ1-38 isotopologue distributions using the centroid values as a measure for average isotope incorporation. For Scheme 1 **(A,B),** statistics on the centroid values (green line: Aβ1-38; red line: Aβ1-42), show no significant difference in isotope incorporation, while in Schemes 2 and 3 the degree of isotope incorporation in plaques was statistically different in Aβ1-38 compared to Aβ1-42 (right: centroid value; left: centroid normalized to number of nitrogen atoms per peptide sequence) **(C)** Single ion images of Aβ1-38 for the different centroid m/z values for the corresponding Aβ1-38 and Aβ1-42 isotope distributions as a measure of differential label incorporation. No difference in ion distribution was observed for Aβ1-38 isotopologues showing 15N incorporation levels corresponding to average labelling degree of either Aβ1-42 or Aβ1-38. **(D-F)** For Scheme 2, Aβ1-42 showed significantly higher label incorporation as compared to Aβ1-38 (E) and while no difference in ion distribution are observed for Aβ1-38 isotopologues at m/z for the respective centroid values, as a consequence of broad overlap of the distribution curves **(F)**. **(G,H)** In Scheme 3, Aβ1-38 showed significantly higher incorporation compared to Aβ1-42 as indicated by the isotopologue distribution curves and centroid values. **(I)** This is further reflected in the single ion maps where no unlabeled Aβ1-38 species were observed as seen for Aβ1-42.

While the isotope patterns of Aβ1-38 and Aβ1-42 could not be discriminated using the long labelling approach in Scheme 1, results for Scheme 2 show that the detected Aβ1-38 showed 43% less isotope incorporation as compared to Aβ1-42.

This suggests that all Aβ1-38 has been secreted and deposited during the washout period (Fig. 4D-F), which is further highlighted by single ion images generated for m/z shifts corresponding to the maximal Aβ1-42 label incorporation detected (Supplemental Information Fig. S11). Further, we performed two additional PULSE/CHASE experiments based on Scheme 2 (4 weeks PULSE, 8 weeks CHASE), with modified CHASE (4 weeks, Scheme 2b, and 10 weeks, Scheme 2c). These data confirm both differential labelling of Aβ1-42 and Aβ1-38 as well as show similar rates of label washout (Supplementary Information Fig. S12) and further support that Aβ1-42 is secreted and deposited prior to Aβ1-38.

This was even more prominent in Scheme 3, with labelling starting at week 10. Here, only labelled Aβ1-38 was detected, which is in contrast to Aβ1-42, that show less than 50% less average label incorporation along with pronounced deposition of unlabeled Aβ1-42. This further suggests that Aβ1-38 secretion and deposition is taking place at a later point of time as compared to Aβ1-42 (Fig. 4G-I). Together, this validates that both production and deposition of Aβ1-38 represent a downstream event that occurs only after the initial seeding and plaque formation through deposition of Aβ1-42.

## Discussion

We employed a novel chemical imaging approach, iSILK, based on metabolic labelling with stable isotopes followed by multimodal imaging mass spectrometry, for delineating the early events in developing plaque pathology in APP^NL-G-F^ mice. This approach allowed us to visualize Aβ aggregation dynamics in different brain regions as well as the chemical basis for development of heterogenic intra-plaque morphology. Using a comprehensive PULSE/CHASE labelling scheme setup, we found that Aβ pathology in APP^NL-G-F^ mice precipitates first in the cortex by formation as a small dense core, with deposits consisting of Aβ1-42. Later events in early plaque pathology involve homogenous Aβ1-42 deposition and plaque growth in the cortex followed by deposition in the hippocampus, and eventually secretion and deposition of Aβ1-38.

While seeding of Aβ pathology is considered a key event in early amyloid plaque formation, the mechanism of subsequent mature amyloid plaque growth is not well understood. Familiar and sporadic cases of AD display structurally distinct plaque pathology. ^21, 34^ Further, the morphological plaque heterogeneity has been suggested to depend on underlying Aβ peptide composition, where presence of the core has been associated with deposition of shorter Aβ peptides in both sporadic and familiar cases. In sporadic AD it has been suggested to represent a secondary mechanism in plaque growth. ^21^ In terms of AD-causing mutations on the other hand, this would be a result of either overproduction of total Aβ (*e.g*., Swedish APP mutation), or increased aggregation propensity of the Aβ peptide itself (*e.g*., Arctic APP mutation) ^30, 31^. Further, a shift towards increased Aβ1-42 production instead of Aβ1-40 caused by the Iberian APP mutation is observed for the APP^NL-G-F^ mice in our study. However, contrary to our results, this shift towards increased Aβ1-42 has not been associated with a distinct formation of amyloid cores as seen here, but rather widely spread diffuse plaque pathology ^23, 35^. In addition, multiple mutations have previously been described to have an effect on plaque morphology, in particular core formation, as demonstrated by studies in transgenic APP mice carrying the Swedish-, or double Arctic x Swedish mutation ^31^. Here, transgenic Swedish mice develop large plaques with single core structure whereas transgenic mice with Arctic x Swedish double mutations develop plaques with multiple small cores within a single plaque ^31^. Therefore, given the clear initial plaque core formation in the APP^NL-G-F^ mice used here, as indicated by the ^15^N/^14^N NanoSIMS data, the high aggregation state of these structures identified by LCO double staining, and the almost exclusive presence of the Aβ1-42, the plaque core formation in these mice represents a cumulative effect of the three mutations.

Further, our data indicate that the Aβ1-38 is the second main peptide present in the APP^NL-G-F^ mice. Recently, MALDI-based 3D-reconstruction of amyloid plaques in aged APP^NL-G-F^ mice suggested the presence of Aβ1-42 in cores that were surrounded by Aβ1-38 in most but not all plaques. ^20^ iSILK imaging provides strong evidence for a sequence of events that can explain the chemical phenotype of these plaques. This is further in line with ELISA results for these mice ^23^, where the other main peptide detected was the Aβ1-38. Interestingly, presence of shorter C- and N-terminally truncated peptides within single plaques has been suggested to represent intra-plaque metabolism of Aβ1-42 towards shorter fragments, a process facilitated by enzymatic activity ^36, 37^. While this might still be the case for the N-terminally truncated fragments as shown by the similar distribution pattern of the Aβx-40 and Aβx-42 ^21^, this is likely not the case for the C-terminally-truncated peptides. Here, given that ^14^N Aβ1-38 was not detected in either feeding Scheme 1 or 3, our data suggest that the generation of Aβ1-38 represents an independent event, probably through γ-secretase-mediated APP processing.

In summary, we show that metabolically labeled APP and Aβ accumulate in plaques upon pathology onset and can be traced with multimodal imaging mass spectrometry at cellular length scales. We identified the cortex as the primary deposition site in precipitating amyloid pathology in the current APP^NL-G-F^ knock-in model of AD. Further, we show that in this model, plaque formation involves an initial deposition of a small, q-FTAA-positive core seed made up exclusively of Aβ1-42. During evolving plaque pathology, small, cored deposits grow outwards, radially, following deposition of later secreted Aβ1-42 and Aβ1-38.

## Conclusion

Taken together, the data presented here show that iSILK is a very promising approach to study the Aβ plaque formation dynamics over time in experimental AD. This in turn can provide novel insights on early amyloid pathology, as illustrated here, that are significant to developing strategies to target AD pathogenesis in time particular with respect to ongoing antibody-based drug-trials that target Aβ aggregation intermediates.

The refined methods for imaging metabolically labelled, endogenous peptides and proteins holds further great potential for investigations on studying other genetic mouse models of AD pathology and delineate the effect of different mutations as well as genetic and pharmacological efforts to modifiy Aβ aggregation propensity. The technology can furthermore be of great potential for studying other disease mechanisms such as for studying tumor development in experimental cancer research.

## Methods

The full experimental procedures are specified in the Supplementary Information. Briefly, male APP knock-in mice (APP^NL-G-F^; n=3/scheme) carrying humanized Aβ sequence, along with Swedish mutation (KM670/671NL) on exon 16, as well as Arctic (E693G) and the Beyreuther/Iberian mutations (I716F) on exon 17 were used in the study. For metabolic labelling, Mice were fed with MouseExpress (^15^N, 98%, Cambridge Isotope Lab) mouse feed, based on either ^15^N or ^14^N spirulina. The different labeling experiments comprised feeding ^15^N spirulina from week 7 for 11 weeks (Scheme 1); as well as from week 6 for 4 weeks followed by a 7-week CHASE period (Scheme 2); and in a third labelling scheme (Scheme 3) from week 10 for 4 weeks followed by a 3-week CHASE period. The mice were sacrificed after week 17. Following brain isolation, one half was snap-frozen in liquid N2 cooled isopentane, while the other was immediately dissected into anatomical regions and immersion-fixed in 4% PFA, and thereafter processed for EM. Sagittal cryosections (12 μm) were covered with 2,5-DHAP (Sigma Aldrich) as a matrix using a TM Sprayer (HTX Technologies) for high spatial resolution (10 μm) MALDI-TOF-IMS analysis in linear positive mode (Bruker RapifleX). Bright-field light microscopy images (Zeiss Axio Observer) were collected to guide subsequent EM analysis. Electromicrographs of semi-thin sections (350 nm) of EM prepared tissue were obtained through annular bright-field STEM analysis (GAIA3). The ^15^N incorporation into individual Aβ plaques was performed using an ion microprobe with Cs+ primary ion beam (NanoSIMS N50L). MALDI-IMS adjacent cryosections, were stained with q- and h-FTAA structural amyloid probes. Hyperspectral, confocal images were acquired on a Zeiss LSM780 in order to verify the localization of Aβ plaque, structural delineation of Aβ polymorphism within plaques. LMD of stained plaques was performed and followed by immunoprecipitation for Aβ enrichment and mass spectrometric characterization and validation of Aβ peptides observed in IMS using MALDI-TOF-MS (Bruker Ultraflextreme) and LC-MS/MS analysis (Thermo QExactive Orbitrap).

## Acknowledgements

We thank Jaana Lundgren at the Pathology Division at Sahlgrenska University Hospital and the staff at Centre for Cellular Imaging (CCI), Core Facilities, The Sahlgrenska Academy at the University of Gothenburg for assistance with the EM preparations.

