## Supplementary Information for "Imaging stable isotope labelling kinetics (iSILK) for following spatial Aβ plaque aggregation dynamics in evolving Alzheimer’s disease pathology"

***Content:***
*Supporting Information Figure S1-S11*
*Extended Experimental Section*

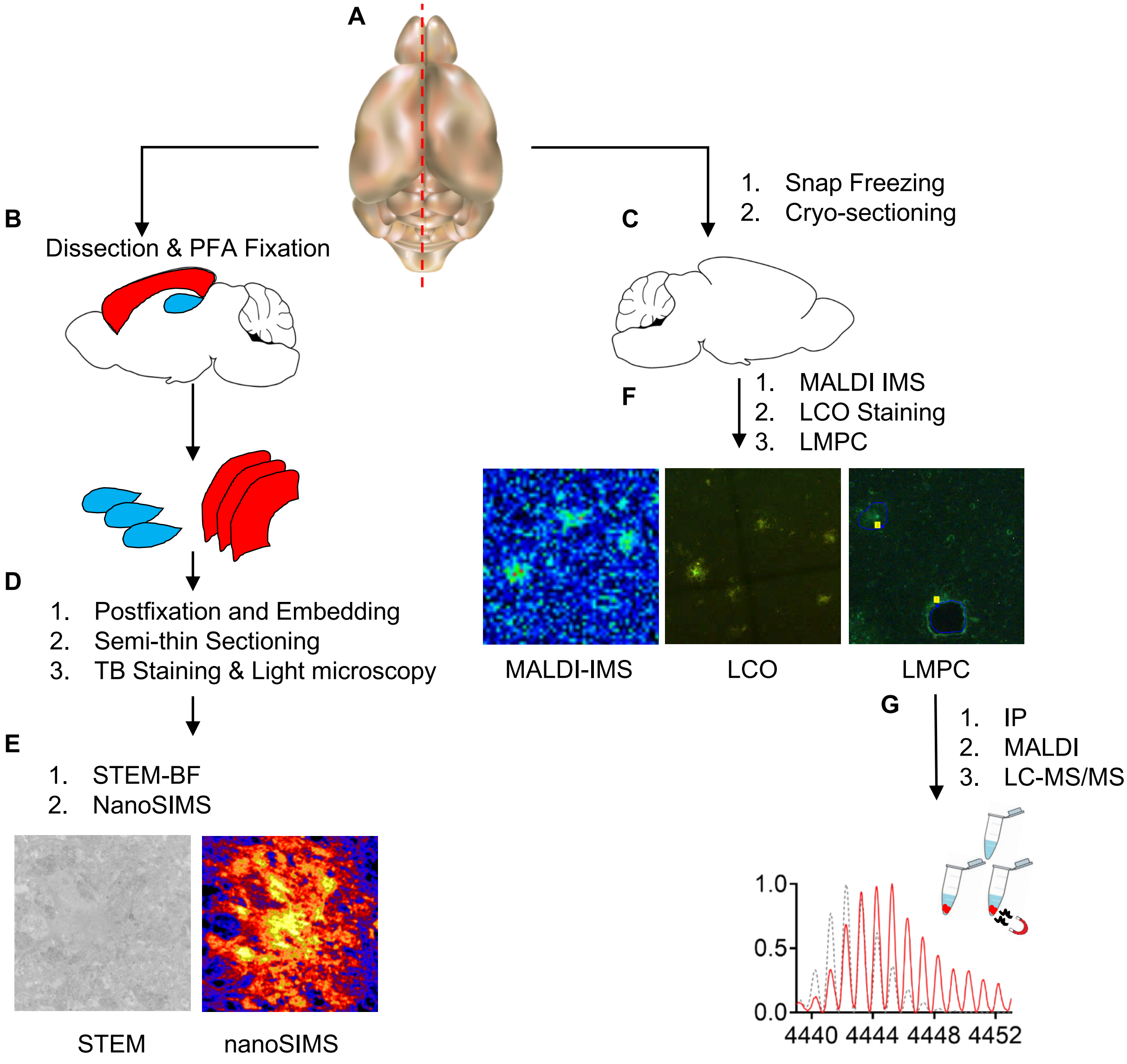


**Supplementary Information Figure S1: iSILK Schematic. (A)** After culling, brains of APP$^{NL\text{-}G\text{-}F}$ mice were isolated and devided sagitally in half. **(B)** One hemisphere was immediately dissected into anatomical reagions and fixed in PFA, while **(C)** other hemisphere was immediately snap frozen for later cryo-sectioning. **(D)** The fixed tissue, was postfixed, embedded for EM, semi-thin sectioned (350µm), and stained with toluidine blue in order to identify smaller ROIs for further analysis. **(E)** Semi-thin sections were collected on formvar coated copper grids and annular STEM-BF images were collected to guide follow-up NanoSIMS analysis. **(F)** Cryo-sectioned samples were analyzed through MALDI-IMS, LCO hyperspectral imaging, and following LMPC, **(G)** through MALDI MS and LC-MS/MS for peptide validation.

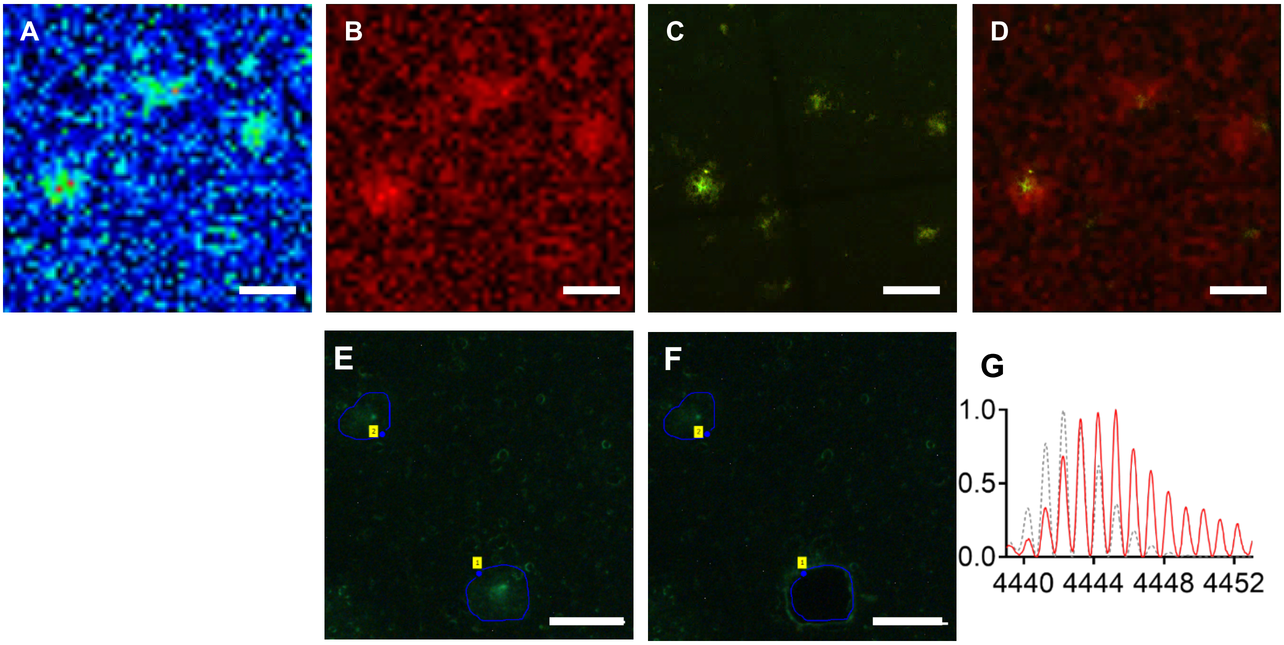


**Supplementary Information Figure S2: MALDI-IMS and LCO-staining on adjacent tissue sections. (A, B)** Sample MALDI-IMS single ion images of the $^{15}N$ Aβ1-42 peptide (A: rainbow scale, B: red scale), show a high degree of colocalization with the **(C)** LCO-based double staining, as seen in **(D)** partial overlay image. Scale bar: A–D: 100 µm;

**(E,F)** LMPC based verification of Aβ peptides. Following LCO double staining, LMPC isolation of individual plaques was performed for downstream MS analysis. Here, **(E)** plaque regions of interest were outlined, and **(F)** laser dissected one-by-one.

**(G)** Immunoprecipitation and MALDI mass spectrometry allowed Aβ peptide identification and validation of isotope incorporation. Scalebar: (A, B) 50 µm.

A

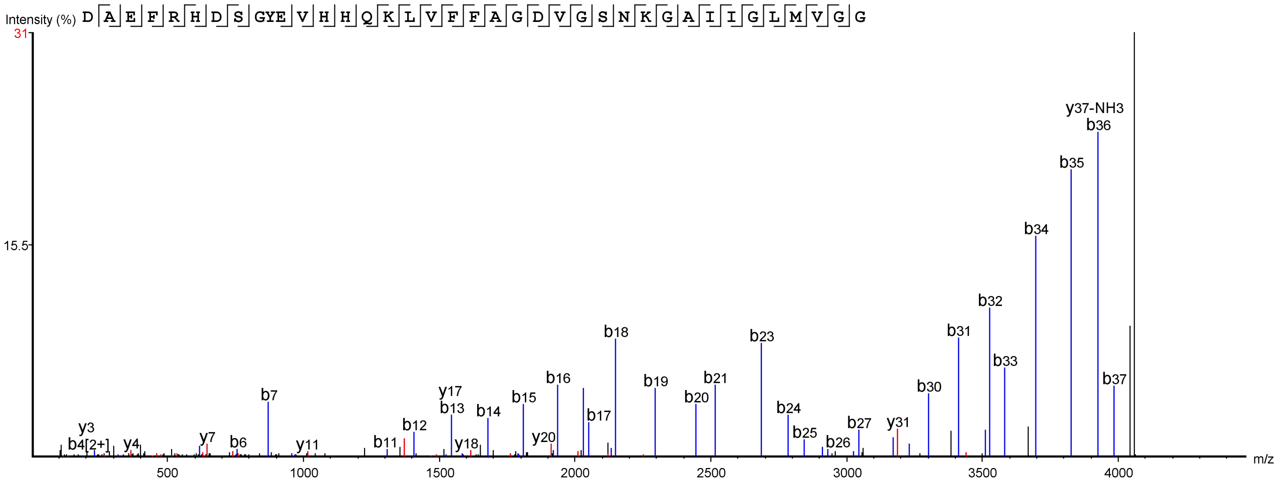


B

**Monoisotopic mass of neutral peptide Mr(calc): 4056.9905**
**Ions Score: 272 Expect: 7.1e-027**
**Matches : 54/196 fragment ions using 80 most intense peaks** (help)

| # | b | b* | $b^0$ | Seq. | y | y* | $y^0$ | # |
|---|---|---|---|---|---|---|---|---|
| 1 | 116.0342 | | 98.0237 | D | | | | 38 |
| 2 | 187.0713 | | 169.0608 | A | 3942.9708 | 3925.9443 | 3924.9602 | 37 |
| 3 | 316.1139 | | 298.1034 | E | 3871.9337 | 3854.9071 | 3853.9231 | 36 |
| 4 | 463.1823 | | 445.1718 | F | 3742.8911 | 3725.8645 | 3724.8805 | 35 |
| 5 | ***619.2835*** | 602.2569 | 601.2729 | R | 3595.8227 | 3578.7961 | 3577.8121 | 34 |
| 6 | ***756.3424*** | 739.3158 | 738.3318 | H | 3439.7216 | 3422.6950 | 3421.7110 | 33 |
| 7 | ***871.3693*** | 854.3428 | 853.3587 | D | 3302.6627 | 3285.6361 | 3284.6521 | 32 |
| 8 | ***958.4013*** | 941.3748 | 940.3908 | S | **3187.6357** | 3170.6092 | 3169.6252 | 31 |
| 9 | 1015.4228 | 998.3962 | 997.4122 | G | 3100.6037 | 3083.5771 | 3082.5931 | 30 |
| 10 | ***1178.4861*** | 1161.4596 | 1160.4756 | Y | 3043.5822 | 3026.5557 | 3025.5717 | 29 |
| 11 | ***1307.5287*** | 1290.5022 | 1289.5182 | E | 2880.5189 | 2863.4923 | 2862.5083 | 28 |
| 12 | ***1406.5971*** | 1389.5706 | 1388.5866 | V | **2751.4763** | 2734.4498 | 2733.4657 | 27 |
| 13 | ***1543.6560*** | 1526.6295 | 1525.6455 | H | **2652.4079** | 2635.3813 | 2634.3973 | 26 |
| 14 | ***1680.7150*** | 1663.6884 | 1662.7044 | H | 2515.3490 | 2498.3224 | 2497.3384 | 25 |
| 15 | ***1808.7735*** | 1791.7470 | 1790.7630 | Q | 2378.2901 | 2361.2635 | 2360.2795 | 24 |
| 16 | ***1936.8685*** | 1919.8420 | 1918.8579 | K | 2250.2315 | 2233.2049 | 2232.2209 | 23 |
| 17 | ***2049.9526*** | 2032.9260 | 2031.9420 | L | 2122.1365 | 2105.1100 | 2104.1260 | 22 |
| 18 | ***2149.0210*** | 2131.9944 | 2131.0104 | V | **2009.0525** | 1992.0259 | 1991.0419 | 21 |
| 19 | ***2296.0894*** | 2279.0628 | 2278.0788 | F | **1909.9840** | 1892.9575 | 1891.9735 | 20 |
| 20 | ***2443.1578*** | 2426.1313 | 2425.1472 | F | **1762.9156** | 1745.8891 | 1744.9051 | 19 |
| 21 | ***2514.1949*** | 2497.1684 | 2496.1844 | A | **1615.8472** | 1598.8207 | 1597.8367 | 18 |
| 22 | ***2571.2164*** | 2554.1898 | 2553.2058 | G | **1544.8101** | 1527.7836 | 1526.7995 | 17 |
| 23 | ***2686.2433*** | 2669.2168 | 2668.2328 | D | **1487.7886** | 1470.7621 | 1469.7781 | 16 |
| 24 | ***2785.3117*** | 2768.2852 | 2767.3012 | V | **1372.7617** | 1355.7351 | 1354.7511 | 15 |
| 25 | ***2842.3332*** | 2825.3067 | 2824.3226 | G | **1273.6933** | 1256.6667 | 1255.6827 | 14 |
| 26 | ***2929.3652*** | 2912.3387 | 2911.3547 | S | 1216.6718 | 1199.6453 | 1198.6613 | 13 |
| 27 | ***3043.4082*** | 3026.3816 | 3025.3976 | N | 1129.6398 | 1112.6132 | | 12 |
| 28 | ***3171.5031*** | 3154.4766 | 3153.4926 | K | **1015.5969** | 998.5703 | | 11 |
| 29 | 3228.5246 | 3211.4980 | 3210.5140 | G | 887.5019 | | | 10 |
| 30 | ***3299.5617*** | 3282.5352 | 3281.5511 | A | 830.4804 | | | 9 |
| 31 | ***3412.6458*** | 3395.6192 | 3394.6352 | I | 759.4433 | | | 8 |
| 32 | ***3525.7298*** | 3508.7033 | 3507.7193 | I | **646.3593** | | | 7 |
| 33 | ***3582.7513*** | 3565.7247 | 3564.7407 | G | **533.2752** | | | 6 |
| 34 | ***3695.8354*** | 3678.8088 | 3677.8248 | L | 476.2537 | | | 5 |
| 35 | ***3826.8758*** | 3809.8493 | 3808.8653 | M | **363.1697** | | | 4 |
| 36 | ***3925.9443*** | 3908.9177 | 3907.9337 | V | 232.1292 | | | 3 |
| 37 | ***3982.9657*** | 3965.9392 | 3964.9552 | G | 133.0608 | | | 2 |
| 38 | | | | G | 76.0393 | | | 1 |

**Supplementary Information Figure S3. LC-MS/MS based verification of Aβ1-38$_{ARCTIC}$.**
**(A)** MS/MS fragment mass spectrum of Aβ1-38$_{ARCTIC}$, where cleavage sites are indicated in the peptide sequence. **(B)** Table of detected and matched fragment ions indicating sequence coverage for Aβ1-38$_{ARCTIC}$. The detected and accurately matched MS/MS fragment ions are color indicated in the spectrum and table (b-ions and y-ions: red).

A

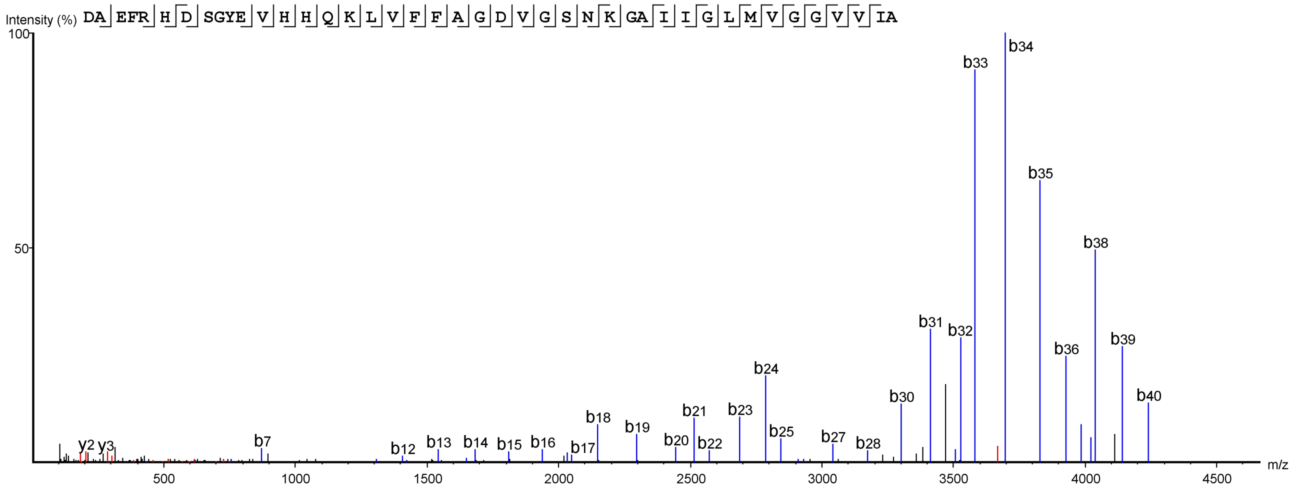


B

**Monoisotopic mass of neutral peptide Mr(calc):** 4439.2485
**Ions Score:** 260 **Expect:** 1.9e-025
**Matches :** 30/212 fragment ions using 41 most intense peaks (help)

| # | b | b* | $b^0$ | Seq. | y | y* | $y^0$ | # |
|---|---|---|---|---|---|---|---|---|
| 1 | 116.0342 | | 98.0237 | D | | | | 42 |
| 2 | 187.0713 | | 169.0608 | A | 4325.2288 | 4308.2023 | 4307.2182 | 41 |
| 3 | 316.1139 | | 298.1034 | E | 4254.1917 | 4237.1651 | 4236.1811 | 40 |
| 4 | 463.1823 | | 445.1718 | F | 4125.1491 | 4108.1226 | 4107.1385 | 39 |
| 5 | 619.2835 | 602.2569 | 601.2729 | R | 3978.0807 | 3961.0541 | 3960.0701 | 38 |
| 6 | 756.3424 | 739.3158 | 738.3318 | H | 3821.9796 | 3804.9530 | 3803.9690 | 37 |
| 7 | 871.3693 | 854.3428 | 853.3587 | D | 3684.9207 | 3667.8941 | 3666.9101 | 36 |
| 8 | 958.4013 | 941.3748 | 940.3908 | S | 3569.8937 | 3552.8672 | 3551.8832 | 35 |
| 9 | 1015.4228 | 998.3962 | 997.4122 | G | 3482.8617 | 3465.8351 | 3464.8511 | 34 |
| 10 | 1178.4861 | 1161.4596 | 1160.4756 | Y | 3425.8402 | 3408.8137 | 3407.8297 | 33 |
| 11 | 1307.5287 | 1290.5022 | 1289.5182 | E | 3262.7769 | 3245.7504 | 3244.7663 | 32 |
| 12 | 1406.5971 | 1389.5706 | 1388.5866 | V | 3133.7343 | 3116.7078 | 3115.7237 | 31 |
| 13 | 1543.6560 | 1526.6295 | 1525.6455 | H | 3034.6659 | 3017.6393 | 3016.6553 | 30 |
| 14 | 1680.7150 | 1663.6884 | 1662.7044 | H | 2897.6070 | 2880.5804 | 2879.5964 | 29 |
| 15 | 1808.7735 | 1791.7470 | 1790.7630 | Q | 2760.5481 | 2743.5215 | 2742.5375 | 28 |
| 16 | 1936.8685 | 1919.8420 | 1918.8579 | K | 2632.4895 | 2615.4629 | 2614.4789 | 27 |
| 17 | 2049.9526 | 2032.9260 | 2031.9420 | L | 2504.3945 | 2487.3680 | 2486.3840 | 26 |
| 18 | 2149.0210 | 2131.9944 | 2131.0104 | V | 2391.3105 | 2374.2839 | 2373.2999 | 25 |
| 19 | 2296.0894 | 2279.0628 | 2278.0788 | F | 2292.2421 | 2275.2155 | 2274.2315 | 24 |
| 20 | 2443.1578 | 2426.1313 | 2425.1472 | F | 2145.1736 | 2128.1471 | 2127.1631 | 23 |
| 21 | 2514.1949 | 2497.1684 | 2496.1844 | A | 1998.1052 | 1981.0787 | 1980.0947 | 22 |
| 22 | 2571.2164 | 2554.1898 | 2553.2058 | G | 1927.0681 | 1910.0416 | 1909.0575 | 21 |
| 23 | 2686.2433 | 2669.2168 | 2668.2328 | D | 1870.0466 | 1853.0201 | 1852.0361 | 20 |
| 24 | 2785.3117 | 2768.2852 | 2767.3012 | V | 1755.0197 | 1737.9932 | 1737.0091 | 19 |
| 25 | 2842.3332 | 2825.3067 | 2824.3226 | G | 1655.9513 | 1638.9247 | 1637.9407 | 18 |
| 26 | 2929.3652 | 2912.3387 | 2911.3547 | S | 1598.9298 | 1581.9033 | 1580.9193 | 17 |
| 27 | 3043.4082 | 3026.3816 | 3025.3976 | N | 1511.8978 | 1494.8712 | | 16 |
| 28 | 3171.5031 | 3154.4766 | 3153.4926 | K | 1397.8549 | 1380.8283 | | 15 |
| 29 | 3228.5246 | 3211.4980 | 3210.5140 | G | 1269.7599 | | | 14 |
| 30 | 3299.5617 | 3282.5352 | 3281.5511 | A | 1212.7384 | | | 13 |
| 31 | 3412.6458 | 3395.6192 | 3394.6352 | I | 1141.7013 | | | 12 |
| 32 | 3525.7298 | 3508.7033 | 3507.7193 | I | 1028.6173 | | | 11 |
| 33 | 3582.7513 | 3565.7247 | 3564.7407 | G | 915.5332 | | | 10 |
| 34 | 3695.8354 | 3678.8088 | 3677.8248 | L | 858.5117 | | | 9 |
| 35 | 3826.8758 | 3809.8493 | 3808.8653 | M | 745.4277 | | | 8 |
| 36 | 3925.9443 | 3908.9177 | 3907.9337 | V | 614.3872 | | | 7 |
| 37 | 3982.9657 | 3965.9392 | 3964.9552 | G | 515.3188 | | | 6 |
| 38 | 4039.9872 | 4022.9606 | 4021.9766 | G | 458.2973 | | | 5 |
| 39 | 4139.0556 | 4122.0290 | 4121.0450 | V | 401.2758 | | | 4 |
| 40 | 4238.1240 | 4221.0975 | 4220.1134 | V | 302.2074 | | | 3 |
| 41 | 4351.2081 | 4334.1815 | 4333.1975 | I | 203.1390 | | | 2 |
| 42 | | | | A | 90.0550 | | | 1 |

**Supplementary Information Figure S4. LC-MS/MS based verification of Aβ1-42$_{ARCTIC}$.**
**(A)** MS/MS fragment mass spectrum of Aβ1-42$_{ARCTIC}$, where cleavage sites are indicated in the peptide sequence. **(B)** Table of detected and matched fragment ions indicating sequence coverage for Aβ1-42$_{ARCTIC}$. The detected and accurately matched MS/MS fragment ions are color indicated in the spectrum and table (b-ions and y-ions: red).

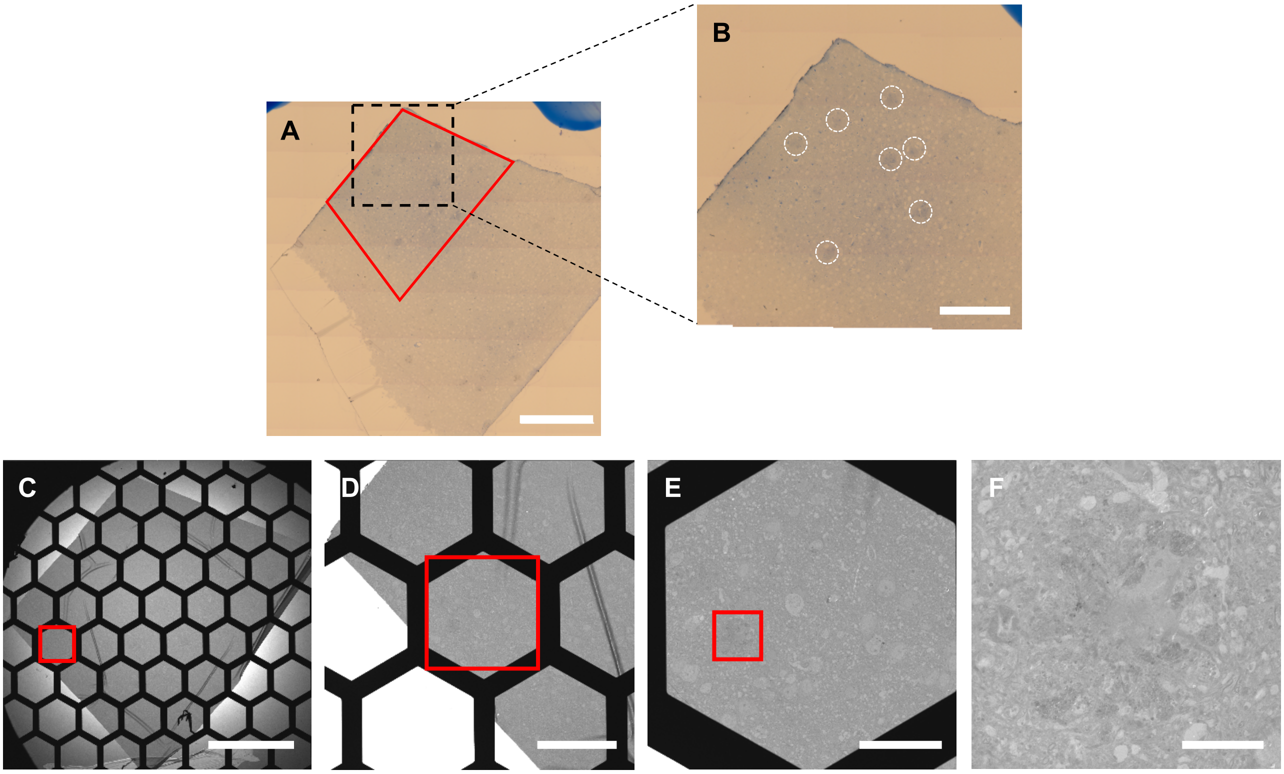


**Supplementary Information Figure S5: Light-microscopy guided STEM-BF imaging of Aβ plaques. (A)** Semi-thin slices were stained with toluidine blue, and bright field light microscopy images were aquired in order to identify plaque-rich areas (red trapezoid) as seen in the **(B)** zoom image. Following aquisition of an **(C)** overview STEM-BF image, the light microscopy images were used to aid in quick identification of plaque. Additional images with **(D, E)** gradually increasing magnification were aquired to aid in downstream localization during NanoSIMS analysis, while keeping the Aβ plaque of interest in focus (red square). **(F)** Finally, high resoltuion images were aquired of individual Aβ plaques. Scalebar: (A,C) 500µm, (B) 300µm, (D) 150µm, (E) 60µm, (F) 10µm.

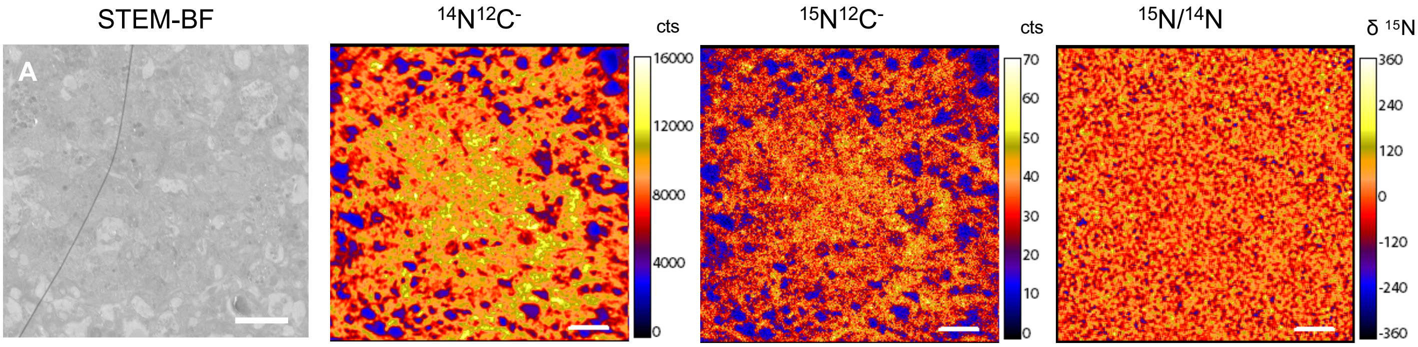


**Supplementary Information Figure S6: STEM-BF and NanoSIMS imaging of Aβ plaques in $^{14}N$-feed control animals.** To assess the natural of $^{15}N$ in the unlabelled NanoSIMS **(A)** STEM- BF images of control unimals (fed with $^{14}N$ protein diet) were aquired. Follow-up NanoSIMS analysis show m/z 26 $^{14}N^{12}C^-$ **(B)** and m/z 27 $^{15}N^{12}C^-$ signal **(C)** to be associated with amyloid plaques. **(D)** Differential (delta) image showing enrichment in permil. Only natural $^{15}N/^{14}N$ content corresponding to 0.367% was present. Scalebar: (A-C) 5μm.

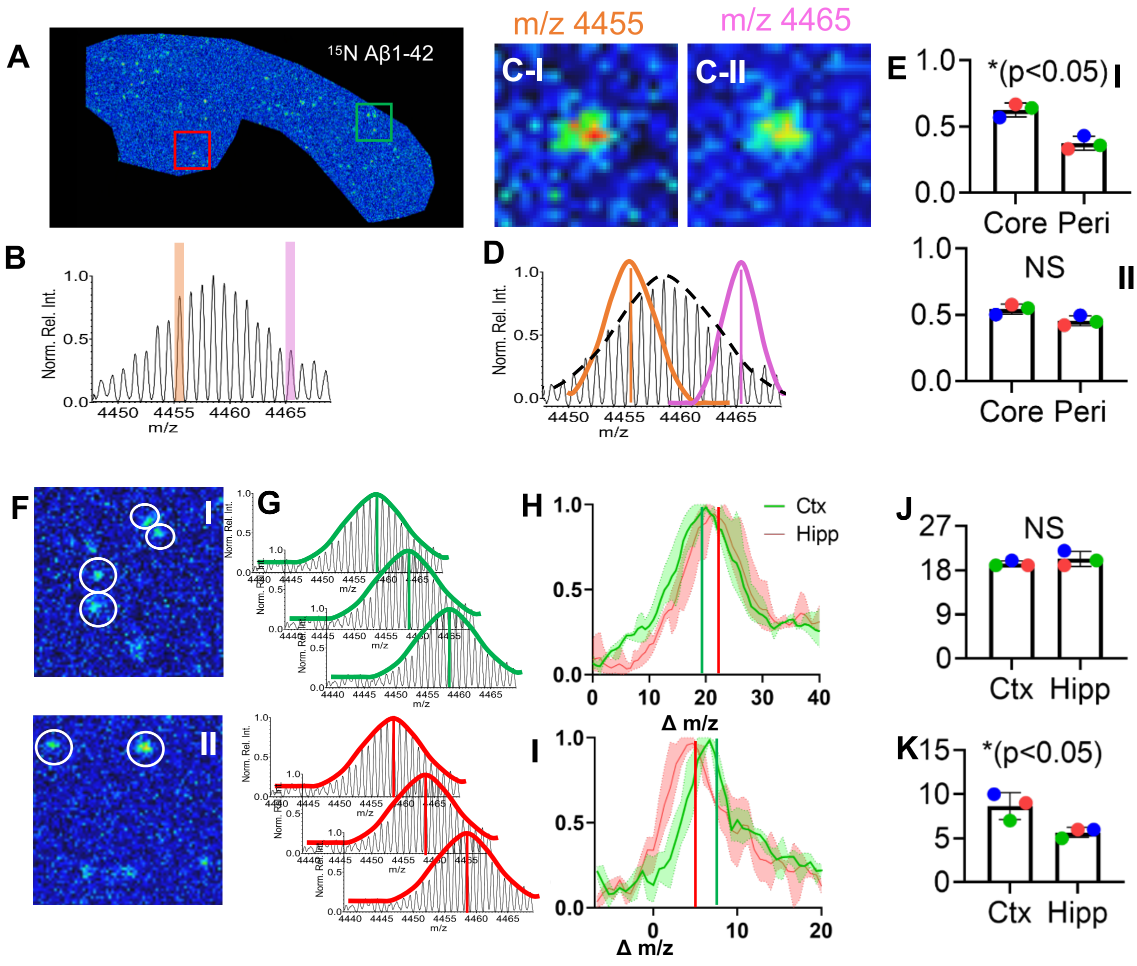


**Supplementary Information Figure S7: Data Analysis Strategy of isotopologue data.**

**(A)** Single ion map of total 15N- Aβ1-42 showing plaques across ctx and hippocampus. **(B) I**sotopic envelope of Aβ1-42 showing different isotopologues. **(C)** Single ion maps for different isotopologues of Aβ1-42 ie isoforms with different degree of $^{15}$N as indicated by the mass shift. Here, less labelled Aβ1-42 (orange, m/z 4455) localizes to the core while more labelled Aβ1-42 (purple, m/z 4465, **C-I**) localized more homogenously across the plaque **(C-II)**.

**(D)** For statistical analysis, mass envelope data were extracted for core (orange) and periphery (purple) ROI of n<10 plaques per animal. The centroid of the distribution curve fitted to the corresponding ROI isotope envelope (core vs periphery) serves as unbiased parameter as mass shift ie. degree of isotope incorporation for statistical analysis. Dotted line illustrates average signal distribution. **(E-I)** Comparative statistics of peak intensities in between core and periphery for the centroid isotopologue of the core (m/z 4455) showing significantly higher levels in the core. **(E-II)** In contrast no significant difference in peak intensity was observed for the centroid isotopologue of the periphery (m/z 4465).

**(F)** Single ion map regions from (A, red,green) of total Aβ1-42 outlining cortical **(F-I)** and hippocampal **(F-II)** plaques. **(G)** For each deposit in each region, a distribution curve is fitted to the isotope envelope of Aβ1-42 for curve analytics (centroid value) to determine average isotope incorporation (green: ctx plaques, red: hippocampus) as illustrated in the schematic. **(H,I)** Average distribution curves of plaque Aβ1-42 isotopologue signals in cortical (green) and hippocampal (plaques) for scheme 1 **(H)** and scheme 2 **(I)** with marked centroid values.

**(J,K)** Statistical analysis show no significant difference in average Aβ1-42 isotope pattern centroid values (isotope incorporation) in between the regions **(J)**, though cortical plaques appear less labelled.

For scheme 2 **(I,K)**, cortical plaques contain more label as compared to hippocampal plaques as statistically validated by analysis of the average Aβ1-42 isotope pattern centroid values representing isotope incorporation.

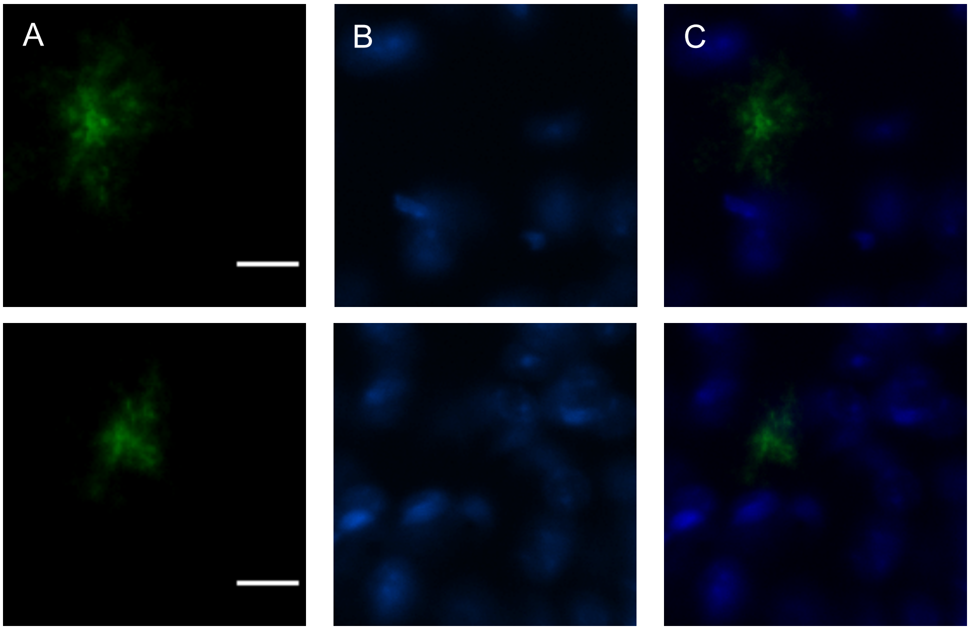


**Supplementary Information Figure S8: Immunohistochemical detection of amyloid plaques in young NLGF mice at 8 weeks. (A)** At time of pathology onset, very small, early plaques (10 µm in size) are detected in the cortex that are with pronounced cores/seeds (ca 5 µm in size) are observed. **(B)** corresponding DAPI staining of nucleii that are of similar size as further isllustrated in the overlay **(C)**. Scalebar: 10 µm

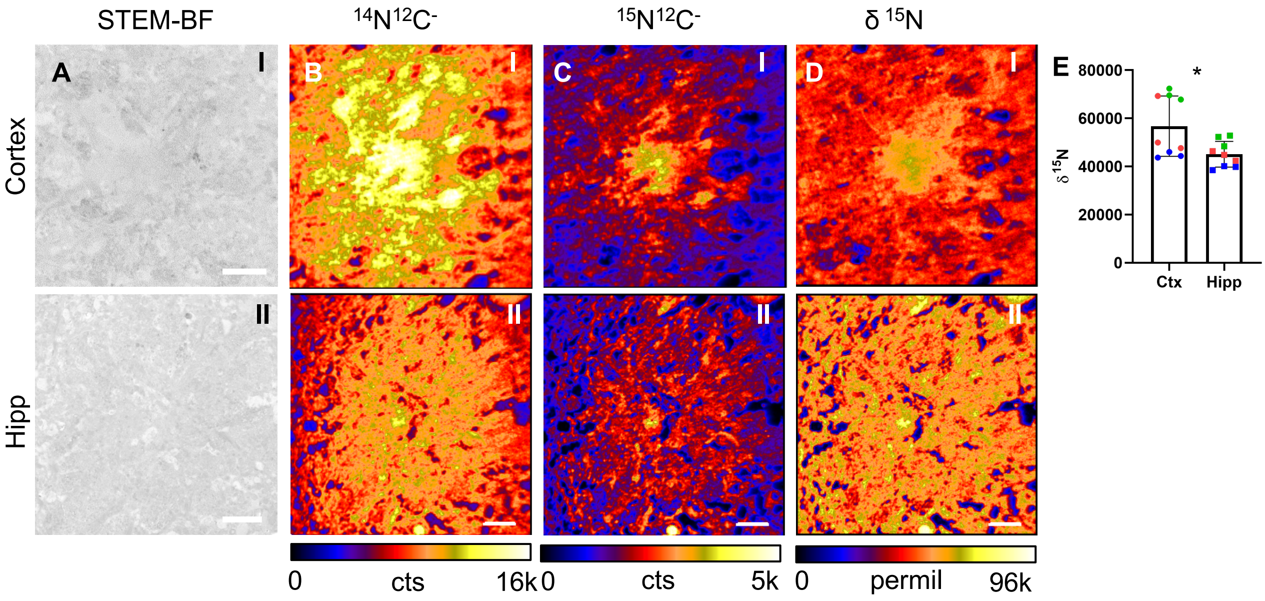


**Supplementary Information Figure S9: NanoSIMS Isotope quantification in single plaques identifies primary regions for plaque deposition. (A)** Sample STEM-BF images of Aβ plaque in cortex (top) and hippocampus (bottom). Corresponding single images for **(B)** m/z 26 $^{14}N^{12}C^{-}$ and **(C)** m/z 27 $^{15}N^{12}C^{-}$. **(D)** Relative $^{15}N$ enrichment in Aβ plaques in both of the anatomical regions. The image shows the deviation of relative $^{15}N/^{14}N$ signal from the natural abundance in permil as calculated from plaque ROI in unlabelled control animals as shown in Fig S6.

**(E)** Comparative analysis of relative enrichment between the two anatomical regions revealed signaficantly more $^{15}N$ enrichment in in cortex, as compared to hippocampus ($p<0.05$) Scalebar: (A-D) 5µm

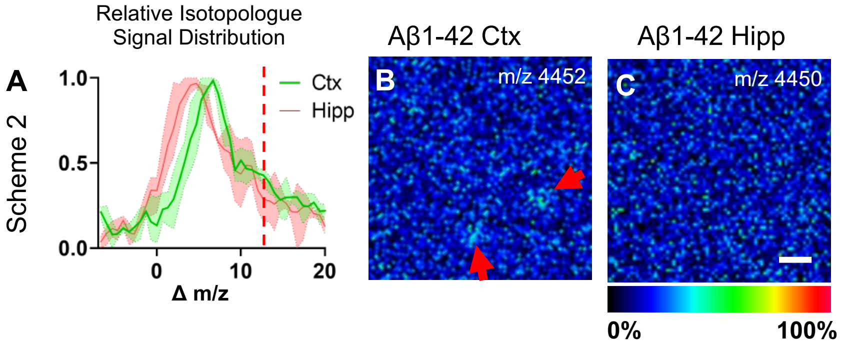


**Supplementary Information Figure S10: Differential localization of Aβ 1-42 species with varying $^{15}N$ incorporation in between cortex and hippocampus. (A)** Average isotopologue distribution curves of Aβ 1-42 signal in cortical –(green) and hippocampal plaques (red). **(B,C)** Single ion maps of $^{15}N$-Aβ 1-42 isotopologue signal corresponding to the maximal mass shift (Δm/z =12; m/z 4452) i.e. degree of isotope inegration for Aβ1-42 observed in the cortex. Here, small plaque features are observed in the cortex (B), while no deposited Aβ1-42 isotopologues at this Δm/z are detected in the hippocampus **(C)**. Scalebar 50 µm

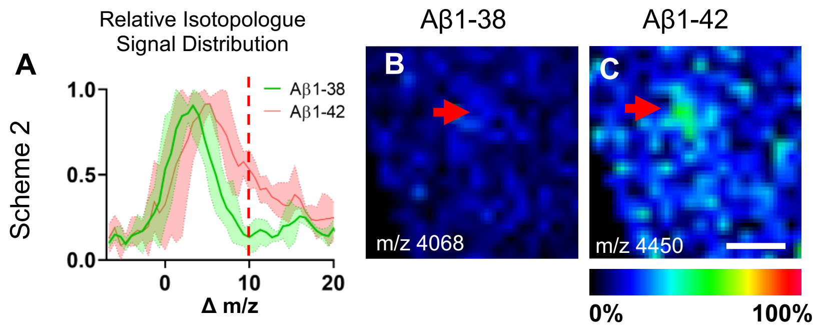


**Supplementary Information Figure S11: Spatial distribution maps of isotopologues for Aβ 1-38 and Aβ1-42.**
**(A)** Ion maps are generated for corresponding isotopologue signal at maximum degree of label incorporation observed for Aβ1-42. No signal of Aβ1-38 is detected **(B)**, in contrast to Aβ1-42, showing distinct accumulation at the core **(C)**. Scalebar: 25 µm

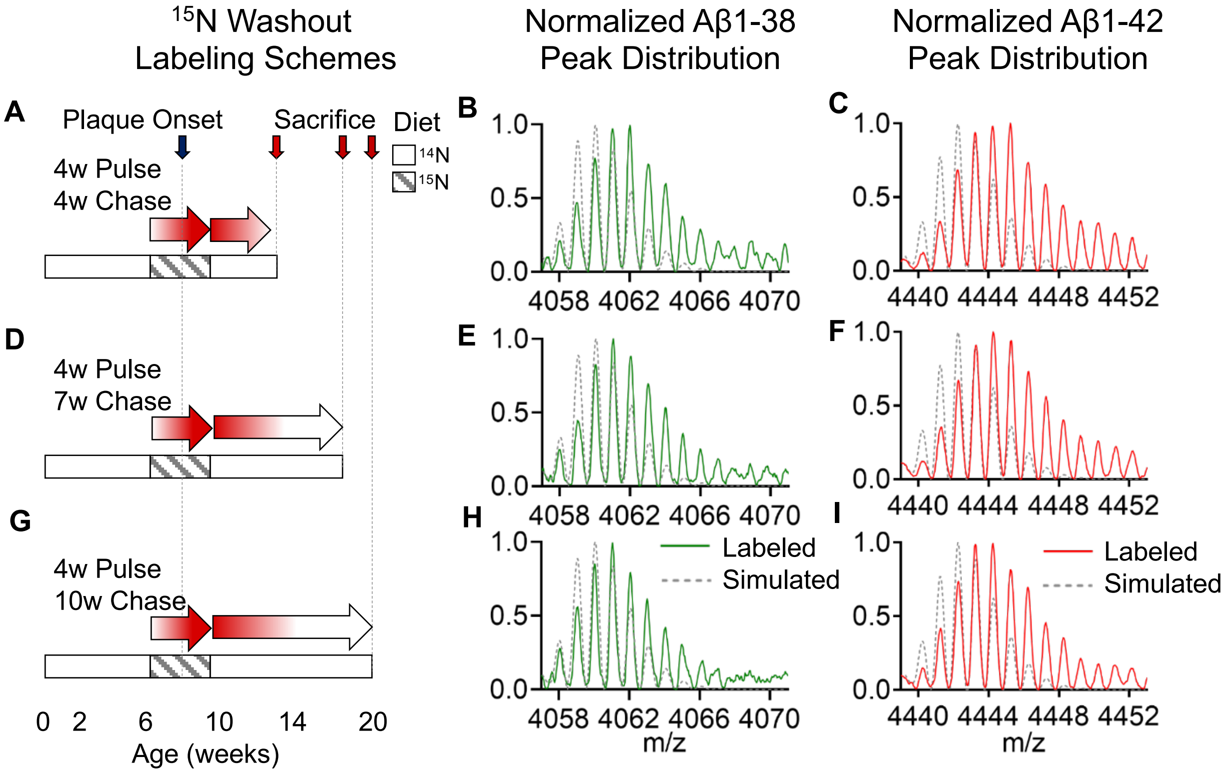


**Supplementary Information Figure S12. Modified CHASE periods show differential labelling and even washout of Aβ1-42 and Aβ1-38.** Additional PULSE(CHASE) experiments based on Scheme 2 **(D)** with different length chase (either 4w **(A-C)**, 7w **(D-F)**, or 10w **(G-I)** following an initial 4week PULSE starting at week 6. MALDI IP-MS data for Aβ1-42 and Aβ1-38 show their maximal degree of $^{15}N$ incorporation at the shortest CHASE time **(B,C)**, which was different in between both species where Aβ1-42 **(C)** showed more label incoporation as compared to Aβ1-38 **(B)**. With extended CHASE (washout), the overall degree of 15N is decreased for both peptides **(E-I)** with Aβ1-42 mainting a higher content of $^{15}N$ as compared to Aβ1-38.

### Experimental Section

#### 1. Chemicals

MouseExpress ($^{15}N$, 98%) mouse feed kit ($^{15}N/^{14}N$) was obtained from Cambridge Isotope Laboratories (Andover, MA, USA). All solvents used in the study were of HPLC/MS grade. Acetone (Ac), acetonitrile (ACN) and absolute ethanol (EtOH), methanol (MeOH) were obtained from Fisher Scientific (Hampton, NH, USA). Glacial Acetic Acid was purchased from VWR Chemicals (Radnor, PA, USA). Paraformaldehyde (PFA), glutaraldehyde, sodium cacodylate buffer, osmium tetroxide and Agar 100 resin were obtained from Agar Scientific (Stansted, Essex, United Kingdom). Formic acid (FA), trifluoroacetic acid (TFA), α-Cyano-4-hydroxycinnamic acid (CHCA), and 2,5-Dihydroxyacetophenone (2,5-DHAP) were obtained from Sigma-Aldrich (St. Louis, MO, USA). TissueTek optimal cutting temperature (OCT) compound was purchased from Sakura Finetek (AJ Alphen aan den Rijn, Netherlands). Indium tin oxide (ITO)-coated conductive glass slides, and peptide calibration standard I were obtained from Bruker Daltonics (Bremen, Germany). PELCO copper grids were obtained from Ted Pella (Redding, CA, USA). The dd$H_2O$ was obtained from a Milli-Q purification system (Merck Millipore, Darmstadt, Germany). The 0.17 polyethylene naphthalate (PEN) membrane slides and Adhesive Cap 500 opaque tubes were purchased from Zeiss/ P.A.L.M. Microlaser Technologies (Bernsried, Germany). Dako Fluorescence Mounting Medium was obtained from Agilent (Santa Clara, CA, USA).

#### 2. Animal Experiment

All procedures and experiments on APP knock in mice (NL-G-F) were performed at UCL together with Prof Frances Edwards in agreement with the Animals (Scientific Procedures) Act 1986, with local ethical approval at UCL (06/05/2016).

Male APP knock-in mice (APPNL-G-F) carrying humanized Aβ sequence, along with Swedish mutation (KM670/671NL) on exon 16, as well as Arctic (E693G) and the Beyreuther/Iberian mutations (I716F) on exon 17 were used in the study. To study spatio-temporal Aβ plaque metabolism, mice were fed MouseExpress (15N, 98%) mouse feed (PULSE), based on $^{15}N$ following the different labelling schemes:
Scheme1: PULSE 10weeks (week 7 to 17); no CHASE (n=3)
Scheme 2 PULSE 4 weeks (week 6 to 10), CHASE 7 week; (n=3)
Scheme 2b PULSE 4 weeks (week 6 to 10), CHASE 4 week, (n=3)
Scheme 2c PULSE 4 weeks (week 6 to 10), CHASE 10 week; (n=3)
Scheme 3: PULSE 4 weeks (week 10-14), CHASE 4 week. (n=3)
Control animals (n=3) for each scheme received 14N spirulina diet.
The mice were sacrificed after the CHASE period. Following brain isolation, one half was snap-frozen in liquid N2 cooled isopentate, while the other was immediately dissected into anatomical regions and immersion-fixed in 4% PFA, and thereafter processed for EM.

#### 3. Tissue Preparation

##### Frozen tissue preparation

One hemisphere was snap frozen directly after isolation using liquid nitrogen (− 150 °C)-cooled isopentane (Supporting Information Fig 1). For MALDI-IMS (and LCO analysis) 12µm cryosections were collected from fresh frozen brain tissue on a cryostat microtome (Leica CM 1520, Leica Biosystems, Nussloch, Germany) at -18°C. For MALDI IMS, the sections were thaw mounted on conductive indium tin oxide (ITO) glass slides (Bruker Daltonics, Bremen, Germany). For LMPC, 12µm thick fresh frozen sections were cut on and mounted on 0.17 PEN membrane slides. All tissue was stored at -80°C.

**EM tissue preparation of PFA fixed tissue**

For EM and NanoSIMS, following harvesting one brain hemisphere was immediately dissected into anatomical regions and immersion-fixed in 4% PFA at 4°C (Suppl. Fig. 1). Tissue was then transferred to modify Karnovsky fixative, containing 2% paraformaldehyde, and 2% glutaraldehyde in 0.1M sodium cacodylate buffer, for 4 hours (1h at RT, 3h at 4°C, on a rotor). Further tissue processing was performed using Automatic Microwave Tissue Processor (Leica EM AMW, Leica Microsystems, Wetzlar, Germany). Here, tissue was washed 4 times with 0.1M sodium cacodylate buffer. Tissue was postfixed with 1% OsO4, and 1% K-Fe in 0.1M sodium cacodylate buffer for 45 min at RT. For additional contrasting, *en bloc* staining with 0.5% Uranyl Acetate (UA) in $ddH_2O$, was performed for 45min. Dehydration was done with rising concentrations of EtOH (50%, 70%, 85%, 95%, and 2 times with absolute EtOH) and 2 times with 100% acetone at RT, 10 minutes each. Samples were embedded in Agar 100 resin, and the resin was polymerized for 24h at 60°C. All samples within an isotopic-labelling scheme were embedded using the same batch of resin, under the same conditions. Semi-thin tissue sections (350nm – diamond knife) were obtained with an ultra-microtome (Leica EM UC6, Leica Microsystems, Wetzlar, Germany). Here, first 350 µm sections were collected on glass microscope slides and stained with toluidine blue for light microscopy, in order to identify Aβ plaque rich regions of the resin block and guide EM analysis. The block was trimmed and 350nm thick serial sections were cut and collected onto formvar coated PELCO copper grids (Suppl. Fig S5).

### 4. EM analysis

Electron microscopy observations were carried out on a Focused Ion-Beam combined with a scanning electron microscope, FIB-SEM workstation through annular bright-field scanning transmission electron microscopy, STEM analysis (GAIA3 Tescan, Brno-Kohoutovice, Czech Republic). The instrument was operated at 30.0 kV, with WD of 5.3mm, yielding a spot size of approx. 7nm, and effective pixel size of approx. 20nm at 12000x magnification used for collection of single plaque images. Image treatment and analysis was performed in Fiji and Adobe Photoshop.

### 5. NanoSIMS data acquisition

Following STEM analysis of fixed samples, ion microprobe (NanoSIMS 50L, Cameca) measurements were performed on the same copper grids. Sections were gold-coated before being transferred to the NanoSIMS.

NanoSIMS images were obtained by rasterizing a 16 keV $Cs^+$ primary ion beam across the sample surface to generate negative secondary ions that were extracted and separated by a magnetic sector. For our study, the NanoSIMS was set up to measure the following secondary ions species: $^{12}C^{15}N^-$, $^{12}C^{14}N^-$, $^{12}C^{13}C^-$, $^{12}C^{12}C^-$, $^{19}F^-$, and $^{32}S^-$. The instrument was tuned to a minimum mass resolving power of >8000 (Cameca definition), enough to resolve isobaric interferences.

Prior to acquisition, the area of analysis was implanted with a high current (~22pA), defocused beam to remove the gold coating, implant $Cs^+$ ion into the surface of the sample and reach a sufficient amount of sputtered secondary ions. Images of surfaces (35 × 35 µm or 40 × 40 µm) were then acquired using constant settings (current of 0.7–0.8pA, spot-size of about 150 nm, dwell time = 5 ms; number of pixels = 256 × 256, 10 layers).

In order to provide the isotopic composition of an unlabeled reference tissue, images were taken of isotopically-unlabeled mice Aβ plaques (prepared in an identical manner) at the start of each day of analysis. The mean $^{12}C^{15}N/^{12}C^{14}N$ ratio for the control images was 0.00367794±0.000028 $2\sigma$ (N=5).

A total of four Aβ plaques (N=4), per region and $^{15}N$ isotopically-labelled animal (n=3) were analyzed. Similarly, six Aβ plaques (N=6), were analyzed in $^{14}N$ spirulina fed APP $^{NL-G-F}$ control mice (n=3).

All the obtained data was dead time corrected and aligned before stacking, using the L'IMAGE software (Dr. Larry Nittler, Carnegie Institution of Washington). Regions of interest (ROIs) were drawn around individual Aβ plaques, using the contour lines on the $^{12}C^{14}N^{-}$ image. These Aβ plaques ROIs were used to quantify the mean enrichment of $^{15}N$ in each region. The nitrogen isotope ratio images were obtained by taking the ratio between the cumulated $^{12}C^{15}N^{-}$ and $^{12}C^{14}N^{-}$ images. The deviation (i.e. enrichment) from the natural abundant $^{15}N$ determined in unlabeled control mice is reported in *permil* using the following parameter:

$$\text{Delta value } \delta^{15}N^{-} = \frac{((^{12}C^{15}N^{-}/^{12}C^{14}N^{-})_{measured} - (^{12}C^{15}N^{-}/^{12}C^{14}N^{-})_{reference})}{(^{12}C^{15}N^{-}/^{12}C^{14}N^{-})_{reference}} \times 1000$$

Univariate comparisons between the groups was performed using paired, two tailed t test ($p < 0.05$) (Supporting Information Fig. S.6 and S9)

### 6. Matrix deposition for MALDI-IMS

Prior to MALDI-IMS sections fresh frozen tissue sections mounted on ITO glasses were thawed and dried in a desiccator for 15min. A series of sequential washes of 100% EtOH (60 s), 70% EtOH (30 s), Carnoy's fluid (6:3:1 EtOH/$CHCl_3$/acetic acid) (110 s), 100% EtOH (15 s), H2O with 0.2% TFA (60 s), and 100% EtOH (15 s) was carried out. For enhanced plaque extraction, tissues was subjected to formic acid vapor for 20 minutes. 2,5-Dihydroxyacetophenone (2,5-DHAP) was used as matrix compound and applied using an HTX TM Sprayer (HTX Technologies LLC, Carrboro, NC, USA). A matrix solution of 15 mg/mL 2,5-DHAP in 70%ACN/2%CH3COOH/2%TFA was sprayed onto the tissue sections using the following instrumental parameters: nitrogen flow (10 psi), spray temperature (75°C), nozzle height (40 mm), eight passes with offsets and rotations, and spray velocity (1000 mm/min), and isocratic flow of 100μL/min using 70% ACN as pushing solvent.

### 7. MALDI-IMS Data Acquisition

MALDI-IMS experiments were performed on a rapifleX MALDI-TOF instrument (Bruker Daltonics). Measurements were performed at 5um spatial resolution, with the laser operating at a frequency of 10 kHz, a laser power of 90%, and 200 shots per pixel. Data were acquired in linear positive mode in the mass range of 2000–20000 *m/z* (mass resolution: 500 at 4000 *m/z*). Pre-acquisition calibration of the system was performed using a combination of peptide calibration standard I, and synthetic Aβ peptides (Aβ1-38, Aβ1-39, Aβ1-40, Aβ1-42, Aβ1-43, Aβ1-44, Aβ1-45, Aβ1-46, Aβ1-47, Aβ1-48), in order to ensure calibration over the entire range of potential Aβ species. Acquisition and subsequent processing were performed in FlexImaging (v5.0, Bruker Daltonics).

### 8. LCO double-staining and Immunohistochemistry

A double-stain strategy with two luminescent conjugated oligothiophenes (LCO), based amyloid probes; tetra- and heptameric formyl-thiophene acetic acids (q- and h-FTAA) was used for structural amyloid analysis. Here, ITO glass mounted, 12μm thick fresh-frozen tissue sections, adjacent to those used for MALDI-IMS were analyzed. Prior to staining the sections were thawed in a desiccator and fixed using absolute EtOH, 70% EtOH, and PBS for 10 min each, and double-stained with q-FTAA and h-FTAA (2.4μM q-FTAA and 0.77μM h-FTAA in PBS) similar to a previously described protocol [1,2]. Sections were incubated for 30 min at RT in the dark, rinsed with PBS, desiccated mounted with Dako fluorescence mounting medium, and stored in dark at 4°C.

For Aβ immunolabeling, 6E10 antibody (1:500, Invitrogen) was incubated overnight at 4 °C, followed by secondary antibody staining with Alexa Fluoro 647 (1:1000, Invitrogen).

### 9. Fluorescent Microscopy

Bright-field images of the toluidine blue stained tissue were acquired using a wide field microscope (Axio Observer Z1, Zeiss, Jena, Germany). The images were acquired with a Plan-Apochromat 10x/0.3 DIC objective.

The hyperspectral imaging was performed using an state-of-art inverted laser scanning confocal microscope (ELYRA PS.1 SIM/PAL-M LSM780, Zeiss, Jena, Germany), equipped with a 32-Channel GaAsP spectral detector, in parallel spectral detection design, enabling simultaneous 34-channel spectral readout in lambda mode. The acquisition was performed using a 35nW, 458nm Argon-laser, with the Plan-Apochromat 20x /0.8 objective. The continuous emission was acquired in the range of 405 to 750nm [1,2]. Linear unmixing, a function within the Zen 2011 (Zeiss) software, was used to differentiate between the q-FTAA and h-FTAA fluorescent signals in the double stained samples, and distinguish between true LCO fluorescence spectrum and unwanted autofluorescence, from for instance lipofuscin.

### 10. LMPC based Aβ plaques isolation and mass spectrometry

Prior to microdissection, the tissue was double-stained with q-FTAA and h-FTAA similarly to double-staining for LCO analysis (Supporting Information Fig. S8), using $ddH_2O$ instead of PBS. The tissue was desiccated but not mounted. Microdissection was done using a LMPC microscope (PALM Microbeam LMPC microscope, Zeiss/ P.A.L.M. Microlaser Technologies, Bernsried, Germany) equipped with a 355 nm pulsed UV-laser. The individual Aβ plaques as visualized with LCO double-stain strategy were collected in Adhesive Cap 500 opaque tubes and stored at -20°C prior to extraction. In total an area of 50 000 $\mu m^2$ of amyloid-positive plaques, was microdissected and collected.

#### Aβ Immunoprecipitation

To the isolated amyloid plaques 50µL of 70% formic acid, with 5mM EDTA was added. The samples were sonicated for 5 minutes, incubated for 1h at 24°C. The samples were then neutralized to pH 7 using 0.5M Tris buffer. Aβ peptides were than purified through immunoprecipitation using Aβ-specific antibodies (antibodies 6E10 and 4G8, Signet Laboratories), coupled to magnetic Dynabeads M-280 Sheep Anti-Mouse (Invitrogen) as described previously [3]. The supernatant was collected and dried through lyophilization.

#### Mass Spectrometry

For MS, the samples were reconstituted in 5 µL 20% ACN/0.1% FA. For MALDI MS of peptide extracts the samples were deposited onto a MALDI target using the seed layer preparation. Here a matrix seed layer (CHCA, 20 mg/mL, 90% Ac, 10% MeOH and 0.005% TFA) was pre-spotted onto the target followed by subsequent co-application of 1 µL sample mixed with 1 µL second matrix solution (CHCA, 15 mg/mL, 50% ACN/0.1% TFA). MALDI MS of plaque extracts analysis was performed on a Ultraflextreme MALDI TOF/TOF instrument (Bruker Daltonics) in reflector positive (RP) mode. Data were collected in the mass range of 500–5500 *m/z* (5000 shots, laser frequency: 1000 Hz, laser focus: medium). External calibration was performed by the deposition of a Peptide Calibrant Mix 1 (Bruker Daltonics) spotted adjacent to the sample spots on the target.

Further, to verify the identity of the observed peptides, an LC-MS/MS analysis, using alkaline mobile phase, of brain was carried out using a Q Exactive quadrupole-orbitrap hybrid mass spectrometer equipped with a heated electrospray ionization source (HESI-II) (Thermo Scientific) and UltiMate 3000 binary pump, column oven, and autosampler (Thermo Scientific), as previously described [4], but with the Q Exactive operated in data dependent mode. Briefly, the resolution settings were 70,000 and target values were $1\times10^6$ both for MS and MS/MS acquisitions. Acquisitions were performed with 1 microscan/acquisition. Precursor isolation width was 3 m/z units and ions were fragmented by so-called higher energy collision induced dissociation (HCD) at a normalized collision energy (NCE) of 25.

### 11. Data Analysis.

MALDI IMS data were aligned with LCO fluorescent images obtained from the same section. Plaque ROI (whole plaque, core, periphery) were annotated based on the co-aligned LCO fluorescent images. Total ion current normalized average spectra of the annotated ROIs were exported as *.csv file.

**MALDI IMS Isotope Pattern analysis**

Isotope pattern analysis was performed in GraphPad Prism (v.7). Here, MALDI IMS ROI spectral data were loaded into Prism and average distribution curves fitted to the individual peptide isotope signal pattern. The centroid was used as measure for average isotope incorporation allowing either signal comparison of the corresponding isotopologue (m/z value) as well as for comparative analysis in between distinct ROI (Plaque Ctx vs Hipp) as well as Peptides (1-42 and 1-38). Univariate comparisons between the groups was done using unpaired, two tailed t test ($p < 0.05$). (Supporting Fig. S7)

For individual peak statistics ROI spectra datafiles were imported into Origin (v. 8.1 OriginLab, Northampton, MA, USA) for peak detection and peak width determination using the implemented peak analyzer function. The determined peak widths serve as bin borders for peak integration were exported as tab delimited text file. The bin borders were used for area under curve peak integration within each bin (peak-bin) of all individual ROI average spectra using an in-house developed R script [5]. Data was log transformed. Univariate comparisons of distinct isotopologue signals (m/z) between the intra plaque regions (Core vs Periphery) was done using unpaired, two tailed t test ($p < 0.05$).

**MS/MS Peptide identification**

For MS/MS analysis spectra were deconvoluted using Mascot Distiller before submission to database search using the Mascot search engine (both Matrix Science) as described previously [6]. The MS/MS spectra were searched toward the a local database containing mutant human APP sequences using the following search parameters: precursor mass ± 15 ppm; fragment mass ± 0.05 Da; no enzyme; no fixed modifications; variable modifications including deamidated (NQ), Glu->pyro-Glu (N-term E), oxidation (M); instrument default.


1 Nystrom, S. *et al.* Evidence for age-dependent in vivo conformational rearrangement within Abeta amyloid deposits. *ACS Chem Biol* **8**, 1128-1133, doi:10.1021/cb4000376 (2013).

2 Michno, W. *et al.* Pyroglutamation of amyloid-betax-42 (Abetax-42) followed by Abeta1-40 deposition underlies plaque polymorphism in progressing Alzheimer's disease pathology. *J Biol Chem*, doi:10.1074/jbc.RA118.006604 (2019).

3 Portelius, E. *et al.* Characterization of amyloid beta peptides in cerebrospinal fluid by an automated immunoprecipitation procedure followed by mass spectrometry. *Journal of proteome research* **6**, 4433-4439, doi:10.1021/pr0703627 (2007).

4 Pannee, J. *et al.* Reference measurement procedure for CSF amyloid beta (Abeta)1-42 and the CSF Abeta1-42 /Abeta1-40 ratio - a cross-validation study against amyloid PET. *J Neurochem* **139**, 651-658, doi:10.1111/jnc.13838 (2016).

5 Hanrieder, J. *et al.* L-DOPA-induced dyskinesia is associated with regional increase of striatal dynorphin peptides as elucidated by imaging mass spectrometry. *Molecular & cellular proteomics : MCP* **10**, M111 009308, doi:10.1074/mcp.M111.009308 (2011).

6 Brinkmalm, G. *et al.* An online nano-LC-ESI-FTICR-MS method for comprehensive characterization of endogenous fragments from amyloid beta and amyloid precursor protein in human and cat cerebrospinal fluid. *Journal of mass spectrometry : JMS* **47**, 591-603, doi:10.1002/jms.2987 (2012).
